# Inferring patterns of purifying, positive and balancing selection in the coppery titi monkey (*Plecturocebus cupreus*) utilizing a well-fit evolutionary baseline model

**DOI:** 10.64898/2026.03.03.709419

**Authors:** Vivak Soni, Cyril J. Versoza, John W. Terbot, Gabriella J. Spatola, Karen L. Bales, Susanne P. Pfeifer, Jeffrey D. Jensen

## Abstract

Despite the coppery titi monkey (*Plecturocebus cupreus*) being a model system for the study of neurodevelopment and behavior, the evolutionary forces shaping observed levels and patterns of genetic variation in the species have remained poorly studied. In order to illuminate the pervasive effects of purifying and background selection, we have fit a distribution of fitness effects of newly arising exonic mutations, utilizing patterns of polymorphism and divergence based on a recently published high-quality genome assembly. To further characterize episodically acting selective processes, we additionally performed the first whole-genome scans for recent positive and balancing selection in this species, reducing false-positive rates by incorporating the demographic history of the population into an evolutionary null model. These scans identified a small number of biomedically-relevant genes with strong statistical support for having experienced recent selective sweeps or long-term balancing selection. In addition, we identified four genomic deletions bearing the signatures of balancing selection. Taken together, this study provides the first insights into patterns of persistent and episodic selective processes in this species.

## INTRODUCTION

Native to the Amazonian forests of Peru and Brazil, the coppery titi monkey (*Plecturocebus cupreus*) is a platyrrhine that inhabits the upper and middle canopy of its forest habitat (Heymann et al. 2021). *P. cupreus* is one of 34 recognized species of the genus *Plecturocebus*, itself one of three titi monkey genera, and is a member of the Western Amazon clade of the *moloch* group that diverged ∼4 million years ago, along with *P. moloch*, *P. brunneus*, *P. dubius*, and *P. caligatus* (Byrne et al. 2016, 2018). Coppery titi monkeys belong to the small number of primate species that are monogamous and pair bonded. Paternal care is common in family units (Mendoza and Mason 1986; Valeggia et al. 1999), which can be constituted by up to three generations of offspring (Kinzey 1997). As such, they have been the focus of much neurobiological research into social behavior, particularly on traits related to monogamy (Lukas and Clutton-Brock 2013), pair-bonding and male parenting (Bales et al. 2007, 2017, 2021; Lau et al. 2024), as well as general cognition (Bales et al. 2017; Lau et al. 2020). Yet, despite this focus on phenotypic traits, relatively little is known regarding the evolutionary forces shaping genomic variation in the species. Specifically, the relative contributions of purifying and background selection in dictating genomic variation as well as the episodic effects of positive and balancing selection, and any association between the action of these processes and well-studied phenotypes in this species, have yet to be characterized.

### Inference of the distribution of fitness eIects: long-term selective dynamics

Given that deleterious mutations make up the vast majority of fitness-impacting mutations, the purging of these variants by purifying selection is expected to be the dominant form of selection shaping genomic variation. Relatedly, the effect of this process on linked sites (i.e., background selection (BGS); Charlesworth et al. 1993) additionally shapes both levels and patterns of genomic diversity in the vicinity of functional regions (see the review of Charlesworth and Jensen 2021). As the efficacy of selection is mediated by the effective population size (*N_e_*), the proportion of constrained sites subject to these processes will generally be expected to be greater in populations with larger *N_e_*. As posited by the Neutral and Nearly Neutral Theories of Molecular Evolution (Kimura 1968, 1983; Ohta 1973), what remains in terms of observed genomic polymorphism, as well as observed divergence, is likely dominated by the action of genetic drift, with the potential for relatively rare and episodic contributions from other forms of selection including positive or balancing selection (and see Jensen et al. 2019).

The characterization of the distribution of selection coefficients associated with new mutations (termed the distribution of fitness effects (DFE)) is therefore crucial when developing evolutionary baseline models for population genomic inference (Comeron 2014, 2017; Ewing and Jensen 2014, 2016; Johri et al. 2022a; Morales-Arce et al. 2022; Howell et al. 2023; Terbot et al. 2023; Soni and Jensen 2025). By summarizing the relative proportion of effectively neutral, weakly deleterious, and strongly deleterious mutations in functional genomic regions, the DFE effectively describes the extent of selective constraint (Eyre-Walker and Keightley 2007; Keightley and Eyre-Walker 2010; Bank et al. 2014).

Because the DFE characterizes long-term selection dynamics, divergence data is often utilized together with polymorphism data to infer the DFE in natural populations. One early DFE inference method was the two-step approach introduced by Keightley and Eyre-Walker (2007). Under this approach, 1) a simple, single population size change demographic model is firstly inferred from putatively neutral synonymous site data, and then 2) the DFE is inferred from non-synonymous sites, conditional on that demographic model (Eyre-Walker and Keightley 2007, 2009; Boyko et al. 2008; Schneider et al. 2011). Although numerous variants of this approach have been developed (e.g., Galtier 2016; Tataru and Bataillon 2020), recent studies have shown that neglecting BGS effects (Johri et al. 2021), as well as mutation and recombination rate heterogeneity (Soni et al. 2024), in these approaches can lead to potentially severe mis-inference of the DFE.

In order to account for these factors, an alternative approach for DFE inference was introduced by Johri et al. (2020), utilizing an approximate Bayesian computation framework and forward-in-time simulations to jointly infer population history and the DFE, whilst incorporating underlying mutation and recombination rate heterogeneity. The benefits of such an approach include accounting for the biasing effects of selection on the inference of population history, as well as harnessing information from multiple aspects of the empirical data (including the site frequency spectrum [SFS], divergence data, and linkage disequilibrium [LD]). However, such joint inference is computationally demanding and has therefore been limited to inferring simple population histories with a small number of parameters. For species with whole-genome polymorphism data and coding-sparse genomes however, one can alternatively infer complex demographic models from neutral regions unlinked to directly selected sites (e.g., via coalescent-based methods), prior to performing DFE inference in functional regions conditional on the inferred population history (Soni and Jensen 2025). While similarly a two-step approach as in early implementations, the advantages of this alternative framework include the ability to estimate a highly complex demographic history, as well as the ability to directly account for BGS effects when performing DFE inference.

### Inference of episodic selective events: recent positive selection and balancing selection

Although genome-wide patterns of genetic variation can generally be explained without invoking more episodic selective processes such as selective sweeps or balancing selection, a number of approaches have been developed for the detection of these processes in localized genomic windows, generally relying on expected changes in patterns of variation at linked sites (see the reviews of Stephan 2019; Charlesworth and Jensen 2021 for sweep dynamics, and Soni and Jensen 2024 for balancing selection). Individual selective sweeps are associated with a skew in the SFS toward rare and high-frequency alleles (Braverman et al. 1995; Simonsen et al. 1995; Fay and Wu 2000), as well as a local reduction in nucleotide diversity around the selected locus (Berry et al. 1991). The severity of the hitchhiking effect is determined by a number of factors, including the strength and age of the beneficial mutation, as well as the level of recombination in the given genomic region. Conversely, balancing selection maintains genetic variability in populations (see the reviews of Fijarczyk and Babik 2015; Bitarello et al. 2023). The genomic signatures of balancing selection will vary depending on the age of the segregating balanced allele, with recent balancing selection (in which the age of the balanced mutation is <0.4*N_e_* generations) generating extended LD due to hitchhiking effects as the mutation increases rapidly in frequency, as well as a relative increase in intermediate frequency alleles (Schierup et al. 2000; and see the reviews of Crisci et al. 2012; Charlesworth and Jensen 2021). The pattern of extended LD dissipates once the selected mutation reaches its equilibrium frequency, as recombination subsequently breaks up linkage (Wiuf et al. 2004; Charlesworth 2006; Pavlidis et al. 2012). However, the skew in the SFS toward intermediate frequency alleles becomes more pronounced as the segregation time of the balanced mutation increases, due to new mutations arising on the balanced haplotype (Soni and Jensen 2024). If the balanced mutation segregates for evolutionary timescales of >4*N_e_* generations (so-called ancient balancing selection), and for periods greater than the divergence time of the species in question, trans-species polymorphisms may be detectable (Klein et al. 1998; Leffler et al. 2013).

The theoretical expectations under a model of a single, recent selective sweep, and of balancing selection once the selected mutation has reached equilibrium frequency, provide the basis of composite likelihood ratio (CLR) tests developed for the detection of these processes. Kim and Stephan (2002) developed the first CLR-based method for the detection of selective sweeps, though subsequent studies have demonstrated that neutral evolutionary processes such as recent population bottlenecks can generate similar genomic signatures and thus result in mis-inference (e.g., Jensen et al. 2005; Charlesworth and Jensen 2022). Building on this method, the SweepFinder software (Nielsen et al. 2005; DeGiorgio et al. 2016) utilizes a null derived from the empirically observed SFS in order to model these neutral processes and partially address the problem of mis-inference. For the detection of long-term balancing selection, Cheng and DeGiorgio (2020) developed a mixed-model approach that combines the expectation of the SFS under balancing selection and under neutrality, thereby modelling the change in the SFS shape as the genomic distance increases from the putatively selected site. This modelling approach underlies a suite of CLR-based methods released under the BalLeRMix software package (Cheng and DeGiorgio 2020, 2022). These methods have been shown to increase power to detect balancing selection as the balanced mutation segregates over longer timescales (>25*N_e_* generations; Soni and Jensen 2024). Similar to SweepFinder2, this approach derives a null from the empirical SFS in order to account for alternative and potentially biasing evolutionary processes.

### DFE inference and genomic scans for recent positive selection and balancing selection in non-human primates

The majority of DFE inference studies in non-human primates have applied variants of the Eyre-Walker and Keightley (2007) approach to exonic data from the great apes (e.g., Castellano et al. 2019; Tataru and Bataillon 2020), whilst others have focused on gene regulatory regions (e.g., Simkin et al. 2014; Anderson et al. 2020; Kuderna et al. 2024). More recent studies have harnessed the two-step framework of Soni and Jensen (2025) to infer the DFE in an outgroup species to the haplorrhine lineage, the aye-aye (Soni et al. 2025a), as well as in a species of biomedical interest, the common marmoset (Soni et al. 2025b). Briefly, the DFEs were inferred via forward-in-time simulation under a demographic model previously inferred for aye-ayes (Terbot et al. 2025, and see the review of Soni et al. 2025c) and common marmosets (Soni et al. 2025d) from non-coding regions that are sufficiently distant from functional genomic regions such that BGS effects will not bias demographic inference. The inferred DFEs in both species were characterized by a greater proportion of deleterious variants relative to that inferred in humans by Johri et al. (2023), consistent with the larger long-term *N_e_* in both aye-ayes and common marmosets relative to humans. As with DFE inference, the majority of genome scans in non-human primates have focused on the great apes (e.g., Enard et al. 2010; Locke et al. 2011; Prüfer et al. 2012; Scally et al. 2012; Bataillon et al. 2015; McManus et al. 2015; Cagan et al. 2016; Munch et al. 2016; Nam et al. 2017; Schmidt et al. 2019), though more recent studies have conducted scans for recent positive selection and balancing selection in the aforementioned aye-ayes (Soni et al. 2025e) and common marmosets (Soni et al. 2025b) as well.

In this study, we characterize the DFE and perform genomic scans for selective sweeps and balancing selection in the coppery titi monkey. This quantification was enabled by a number of recent genomic developments in the species. Firstly, the publication of a highly contiguous genome assembly, with gene-level annotations (Pfeifer et al. 2024), allowed for the necessary demarcation of neutral and functional genomic regions. Secondly, the generation of high-quality, whole-genome polymorphism data allowed for an accurate quantification of genome-wide variation in the species. Based on this data, recent studies have provided the basis for an appropriate genomic null model for selection inference, in quantifying the population history of the coppery titi monkey (Terbot et al. 2026), as well as underlying rates and patterns of mutation and recombination (Soni, Versoza et al. 2026; Versoza et al. 2026a, 2026b). Our results suggest that the DFE in coppery titi monkeys is characterized by an excess of effectively neutral mutations relative to humans and common marmosets, another platyrrhine. Power analyses demonstrate that there is limited power to detect episodic forms of selection in coppery titi monkeys, due to the recent and severe population bottleneck experienced by the sampled population. This dynamic is borne out in our genomic scans for selective sweeps and balancing selection, which reveal a small number of candidate genes; however, a number of these putatively targeted genes are biomedically relevant to the coppery titi monkey, given its role as a model system for neurodevelopment.

## RESULTS AND DISCUSSION

### Inferring the DFE in coppery titi monkeys from polymorphism and divergence data

To infer the DFE of new mutation in the coppery titi monkey, we first sought to characterize patterns of exonic divergence between coppery titi monkeys and humans. As information from polymorphism data is limited to relatively short evolutionary timescales (on the order of 4*N_e_* generations), divergence is an important summary statistic to consider when performing DFE inference. To that end, we updated the coppery titi monkey genome in the 447-way mammalian multiple species alignment (Zoonomia Consortium 2020) with the high quality, chromosome-level genome of Pfeifer et al. (2024), and retrieved substitutions between coppery titi monkeys and humans (see the “Materials and Methods” section for details). We found that no individual exon exhibited a higher divergence rate than the maximum neutral divergence observed in non-coding regions by Soni, Versoza et al. (2026) in 1kb windows (Supplementary Figure S1). This result is consistent with expectations, given that purifying selection depresses divergence in functional genomic regions (Charlesworth et al. 1993).

Based on empirical divergence and polymorphism data, we ran forward-in-time simulations in SLiM (Haller et al. 2026) under the coppery titi monkey demographic model inferred by Terbot et al. (2026), illustrated in Figure 1a in order to fit exonic divergence, Watterson’s θ_w_ (Watterson 1975), Tajima’s *D* (Tajima 1989), and the number of singletons per site with a DFE shape comprised of neutral, weakly/moderately deleterious, and strongly deleterious mutational classes. Thus, following a 10*N_ancestral_* generation burn-in phase (where *N_ancestral_* is the initial coppery titi monkey population size), a divergence phase between *P. cupreus* and humans of 32 million years (Glazko and Nei 2003), and a generation time of 6 years (Pacifici et al. 2013; Perez et al. 2013), data was simulated concluding with the demographic model of Terbot et al. (2026). To infer a well-fitting DFE, we began from DFEs previously inferred using a similar approach in humans (Johri et al. 2023) and common marmosets (Soni et al. 2025b), and performed manual grid step iterations from these DFEs based upon the fit to the data (see Figure 1b for the best-fitting DFE, and Figures 1c-f for the fits of the resulting summary statistics to the empirical data). Although the coppery titi monkey is more closely related to the common marmoset than to humans, the inferred DFE exhibits greater similarity to the latter. This is likely owing to the similar long-term *N_e_s* of the coppery titi monkey and humans relative to the much larger *N_e_* in the common marmoset. Specifically, a lower *N_e_* is expected to reduce the efficacy of purifying selection given that the strength of selection acting on an individual mutation is the product of the selection coefficient, *s*, and *N_e_*. Thus, purifying selection would be expected to be less efficient in coppery titi monkeys and humans relative to common marmosets — an expectation consistent with the inferred DFEs from each species.

**Figure 1:**
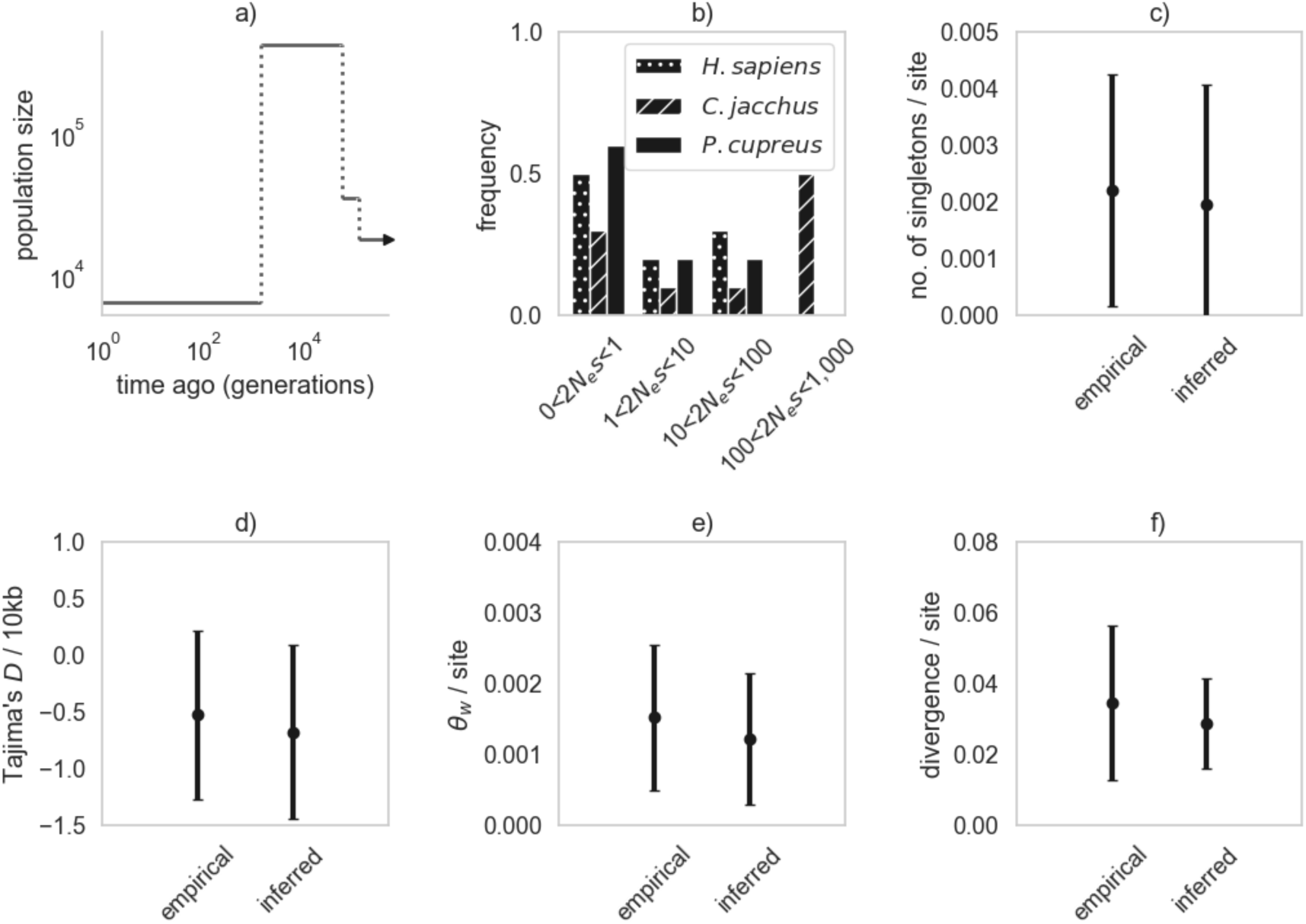
DFE model-fitting results and summaries. **a)** The demographic history previously inferred for this population (re-drawn from Terbot et al. 2026). **b)** Comparison between the best-fitting discrete DFE in the coppery titi monkey inferred in this study (black filled bars), the DFE inferred in another platyrrhine, the common marmoset, by Soni et al. (2025b) (striped bars), and the DFE in humans inferred by Johri et al. (2023) (dotted bars). Exonic mutations were drawn from a DFE comprised of three fixed classes: nearly neutral mutations (i.e., 2*N_ancestral_ s* < 10), weakly/moderately deleterious mutations (10 ≤ 2*N_ancestral_ s* < 100), and strongly deleterious mutations (100 ≤ 2*N_ancestral_ s*). **c–f)** Comparisons of four summary statistics—the number of singletons per site, Tajima’s *D* (Tajima 1989) per 10 kb, Watterson’s *θ*_w_ (Watterson 1975) per site, and exonic divergence per site—between the empirical and simulated data under the inferred DFE. Data points represent the mean value, whilst confidence intervals represent standard deviations.

### Quantifying the power to detect recent positive selection and balancing selection in coppery titi monkeys

A number of factors impact the statistical power to detect positive and balancing selection, including the action of other neutral and selective processes, as well as the nature of the data itself (e.g., Johri et al. 2021, 2022a; Soni et al. 2023, 2024; Soni and Jensen 2024, 2025). It is therefore important to quantify the power to detect episodic processes such as recent positive selection and balancing selection within the context of the demographic history of the population in question, the effects of purifying selection and BGS, and patterns of mutation and recombination rate heterogeneity. We therefore simulated the well-fitting demographic model inferred in coppery titi monkeys by Terbot et al. (2026), using the forward-in-time simulator SLiM (Haller et al. 2026). In order to model the effects of purifying selection and BGS, we utilized the DFE inferred in this study, whilst mutation rates were drawn from the empirical fine-scale mutation rate distribution inferred by Soni, Versoza et al. (2026), and recombination rates were drawn from a normal distribution for each 1kb window, such that the mean rate across each simulation replicate was equal to the mean genomic rate (Versoza et al. 2026b; and see the “Materials and Methods” section for further details).

Figure 2 depicts receiver operating characteristic (ROC) plots across 100 simulated replicates, for selective sweep inference conducted using SweepFinder2 (DeGiorgio et al. 2016), as applied to simulations utilizing two different strengths of selection (2*N_e_s* = 100 and 1,000), and four different times since the introduction of the beneficial mutation (*τ* = 0.1, 0.2, 0.5 and 1.0 *N* generations). Notably, it is only at the very large window size of 100kb that there is reasonable power to detect selective sweeps, and only at the stronger selection value of 2*N_e_s* = 1,000; at smaller window sizes there is little power (see Supplementary Figure S2), owing to the limited variation observed. These results relate to the demographic history of coppery titi monkeys, in which the population has been inferred to experience a severe population size collapse ∼0.08*N_ancestral_* generations prior to sampling, resulting in a current population size ∼1.5% of the size pre-bottleneck. Furthermore, only extremely strong selective sweeps would be expected to be identifiable on this bottlenecked genetic background, thereby reducing inference power, as has previously been described (Barton 1998; Jensen et al. 2005; Thornton and Jensen 2007; Poh et al. 2014; Harris and Jensen 2020).

**Figure 2:**
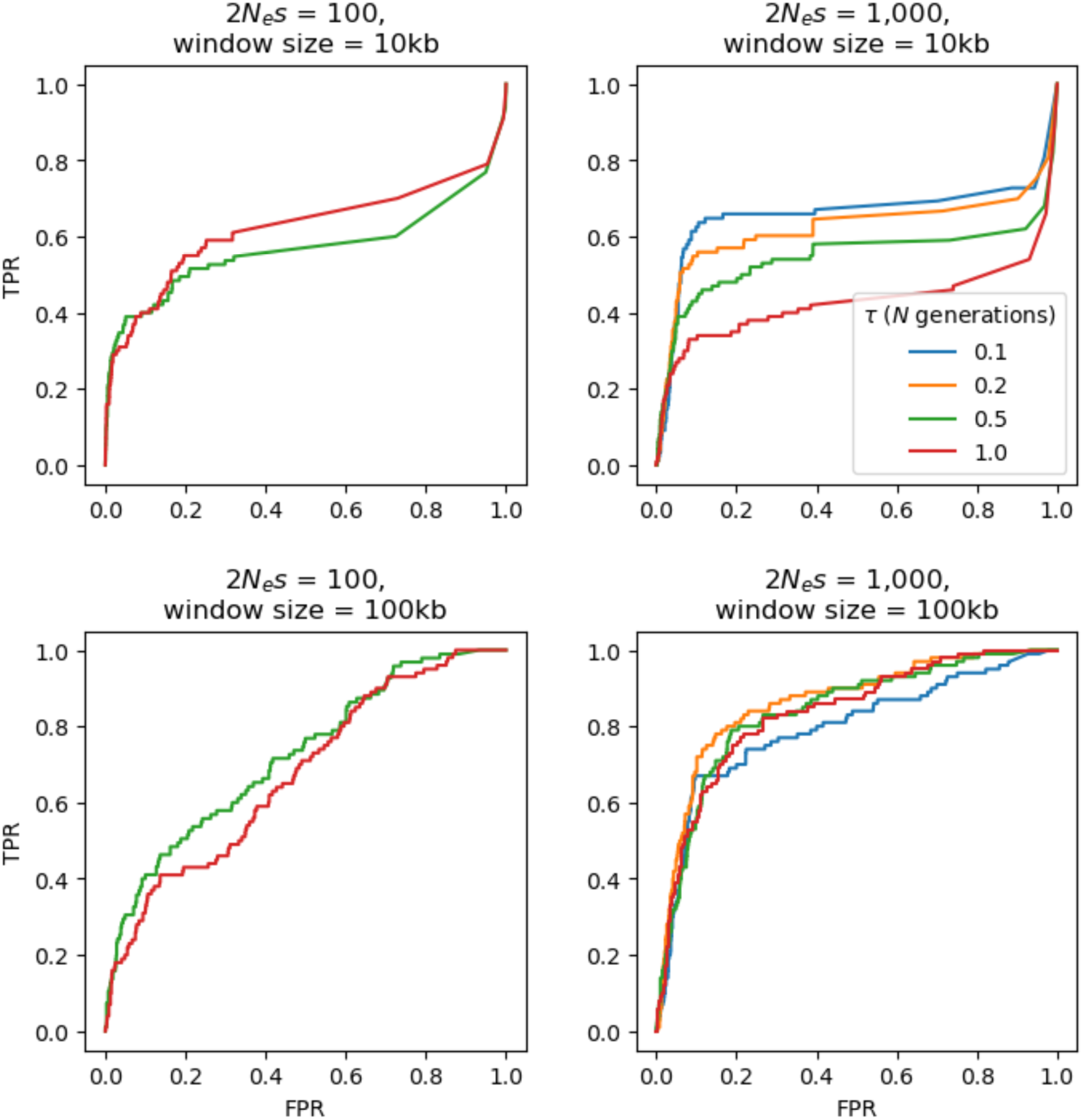
ROC plots based on 100 simulated replicates under the coppery titi monkey demographic model inferred by Terbot et al. (2026) **for selective sweep inference using the SweepFinder2 method.** The false-positive rate (FPR) and the true-positive rate (TPR) are provided on the *x*-axis and *y*-axis of the ROC plots, respectively. Power analyses were conducted across two selection regimes — population-scaled strengths of selection of 2*N_e_s* = 100 and 1000; four times of introduction of the beneficial mutation (*1* = 0.1, 0.2, 0.5, and 1.0 *N* generations ago), and window sizes of 10kb and 100kb. Note that no ROC plots could be plotted for simulated scenarios in which no beneficial mutations reached fixation at the time of sampling.

To quantify the statistical power for the detection of balancing selection using the *B_0MAF_* method implemented in the BalLeRMix package (Cheng and DeGiorgio 2020) under our coppery titi monkey evolutionary baseline model, we introduced a mutation experiencing negative frequency-dependent selection at three different times prior to sampling (*1_B_* = 10, 50 and 75 *N* generations). As shown in Figure 3, power increases with *1_B_*; however, power remains relatively low regardless of window size, even when the balanced mutation has been segregating for extremely long evolutionary timescales. As severe population size reductions will similarly reduce the power to detect balancing selection (Soni and Jensen 2024), this result is consistent with expectation.

**Figure 3:**
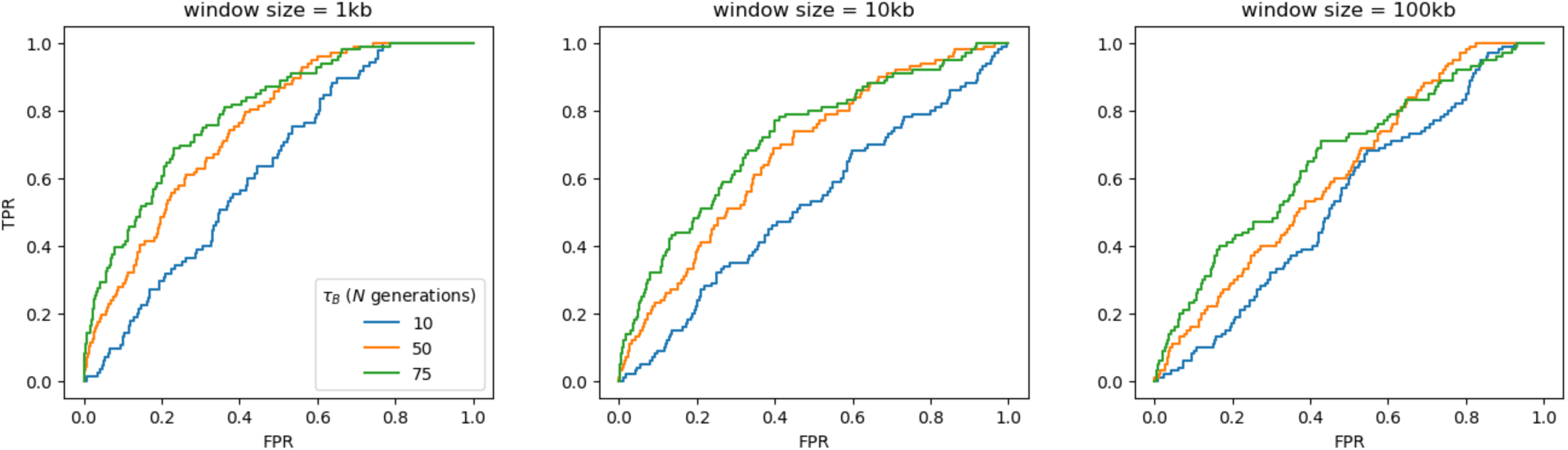
ROC plots based on 100 simulated replicates under the coppery titi monkey demographic model inferred by Terbot et al. (2026) for balancing selection inference using the *B_0MAF_* method. The false-positive rate (FPR) and the true-positive rate (TPR) are provided on the x-axis and y-axis of the ROC plots, respectively. Power analyses were conducted across three times of introduction of the balanced mutation (*1_B_* = 10, 50 and 75 *N* generations), and window sizes of 1kb, 10kb, and 100kb.

Taken together, these power analyses demonstrate the limitations present when attempting to infer these episodic selective processes in recently bottlenecked populations, such as the coppery titi monkey. Moreover, these results highlight the potential perils of more commonly utilized outlier approaches, whereby an arbitrarily chosen threshold is set (e.g., assuming the 1% or 5% tail of loci in the empirical distribution are selection candidates), which have been demonstrated to result in often exorbitant false-positive rates (Teshima et al. 2006; Thornton and Jensen 2007; Jensen et al. 2008; Jensen 2023; Soni et al. 2023). Instead, following the recommendations of Johri et al. (2022a, 2022b), we constructed an evolutionarily appropriate baseline model accounting for commonly operating evolutionary processes that have been previously quantified in coppery titi monkeys. By performing inference with SweepFinder2 (DeGiorgio et al. 2016) and *B_0MAF_* (Cheng and DeGiorgio 2020) on simulated replicates incorporating these species-specific details, we quantified the highest inferred CLR values observed in the absence of positive or balancing selection, and utilized those values as empirical null thresholds (112.3 and 113.8 for the low-mutation and high-mutation rate demographic models in Sweepfinder2). This in turn resulted in ∼4,000 candidates (∼0.04% of all tested loci), when performing inference at each single nucleotide polymorphism (SNP) (Figure 4, and see Supplementary Figures S3-S24 for chromosome-specific plots). The inferred null thresholds for *B_0MAF_* were 236.6 for the low mutation rate model and 280.2 for the high mutation rate model, yielding 3,080 (0.15% of all tested loci) and 1,954 (0.09% of all tested loci) candidates respectively, with inference performed at every fifth SNP (Figure 5, and see Supplementary Figures S3-S24 for chromosome-specific plots). For comparison, a naïve outlier approach with a 5% tail threshold would result in 528,301 sweep candidate loci and 105,661 balancing selection candidate loci, whilst a 1% threshold would yield 105,660 sweep candidates and 21,132 balancing selection candidates. However, as demonstrated in our simulations, the vast majority of these ‘outlier loci’ residing in the tails of the distribution are fully consistent with the baseline model alone, and thus need not invoke positive or balancing selection in order to explain the data (and see Jensen 2023).

**Figure 4:**
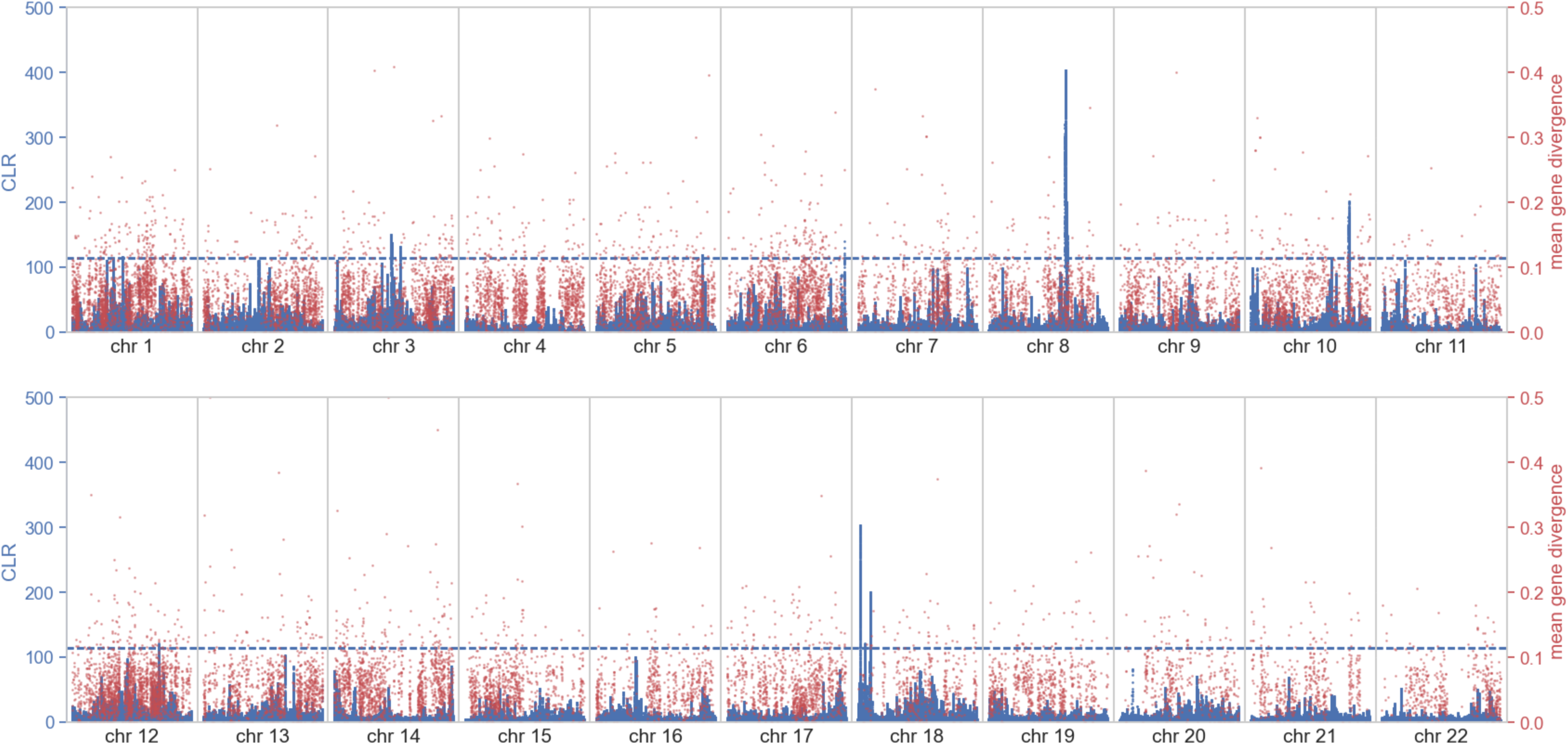
Genome-wide scans for recent positive selection using SweepFinder2 (shown in blue) relative to empirical exonic divergence (red). The x-axis shows the position along the chromosome, the left y-axis shows the composite likelihood ratio (CLR) value of the sweep statistic at each SNP, and the right y-axis provides the mean gene divergence. The horizontal blue dashed line represents the null threshold for sweep detection (note that the null thresholds for the low and high mutation rate demographic models are extremely similar — 112.3 and 113.8 respectively — and are therefore indistinguishable in this figure).

**Figure 5:**
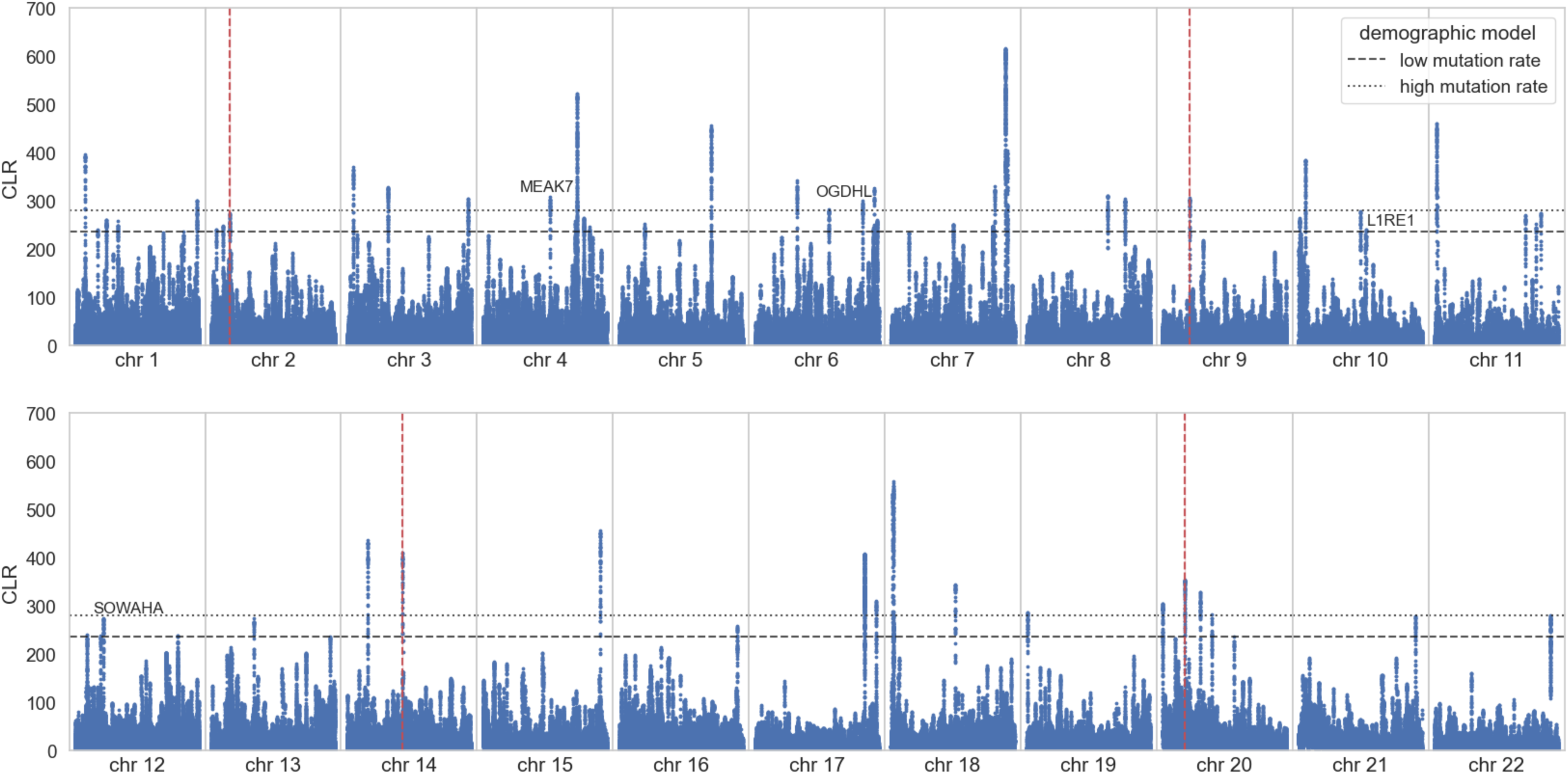
Genome-wide scans for balancing selection using *B_0MAF_*. The x-axis shows the position along the chromosome, and the y-axis the composite likelihood ratio (CLR) value of the balancing selection statistic at each fifth SNP. The horizontal dashed lines represent the null thresholds for the detection of balancing selection, based on two different coppery titi demographic models (see the “Materials and Methods” section for details). The positions of statistically significant deletions are marked by vertical dashed red lines.

### Genomic signatures of positive selection in coppery titi monkeys

The identified selective sweep candidate loci mapped to 47 genes (see Supplementary File S1 for max CLR values per candidate gene). Of these, only 19 were considered well-annotated, based on the percent sequence identity, conserved domains and synteny with humans (see the “Materials and Methods” section for further details). Performing gene functional analysis of these 19 candidate genes using DAVID (Ma et al. 2023) yielded no significant GO terms (see Supplementary Table S1), nor did analysis based upon the most rapidly diverging genes (see Supplementary Table S2).

The gene with the highest CLR values was *PTPN2* (CLR=388.6), identified as a cancer immunotherapy target by Manguso et al. (2017), and involved in the regulation of developmental differentiation of immune cells (Song et al. 2022). Although no previous signatures of positive selection in *PTPN2* have been described in primates, Bello et al. (2023) identified this gene as a selective sweep candidate in an avian species. Given that coppery titi monkeys are a model for neurological development, a notable selective sweep candidate is the gene *MDN1*, a susceptibility gene for epilepsy (Wen et al. 2025), and a previously described sweep candidate in cattle, hypothesized to be related to domestication (Velayudhan et al. 2023). The candidate gene *TMPRSS11D* is also of interest due to high expression levels of this gene in human airways, activating the SARS-CoV-2 spike protein and influenza viruses, a notable result given hypothesized susceptibility amongst non-human primates (Melin et al. 2020); this gene has additionally been recently described as experiencing a hard selective sweep in African populations of humans (Cross et al. 2025).

The candidate gene *ALDH2* plays a key role in the second step of ethanol metabolism, by converting acetaldehyde into acetate (Rodriguez-Zavala et al. 2019). Accumulation of high levels of acetaldehyde have been shown to be both carcinogenic and mutagenic, having adverse effects on the immune system (Ceni et al. 2014; Chen et al. 2014; Zhou et al. 2016; Lee et al. 2017). *ALDH2* has been described as experiencing positive selection in humans (Schaschl et al. 2022), and one may hypothesize that the frugivorous lifestyle of *P. cupreus* may potentially result in the consumption of fermented fruit. Several genome-wide association studies have shown that the genomic region around *ALDH2* is associated with multiple human diseases including rheumatoid arthritis (Coenen et al. 2009), systemic lupus erythematosus (Bentham et al. 2015), type 1 diabetes (Auberger et al. 2014), hypertension (Yasukochi et al. 2017), and coronary artery disease (Wild et al. 2017). This region is therefore of considerable biomedical interest.

### Genomic signatures of balancing selection in coppery titi monkeys

Although the candidate regions from the balancing selection scan mapped to 20 genes (see Supplementary File S1 for max CLR values per candidate gene), only four were considered well-annotated based on the percent sequence identity, conserved domains, and synteny with humans (see the “Materials and Methods” section for further details). Of these, two met the CLR null thresholds for both the low and high mutation rate demographic models. Of the four candidate genes, both *MEAK7* and *SOWAHA* have been identified as selective sweep candidates in other species (Ben-Jemaa et al. 2020), with *MEAK7* being primarily involved in cell proliferation and migration (Nguyen et al. 2018) (note that this gene is also overexpressed in numerous forms of cancer; Nguyen et al. 2019; Chang et al. 2024), whilst *SOWAHA* is a biomarker for colorectal and epithelial cancers (Yi et al. 2024; Yin et al. 2024).

In the context of the coppery titi monkey’s role as a model for neurodevelopment, the candidate gene *OGDHL* is noteworthy. Biallelic variants in *OGDHL* have been proposed to be associated with a number of neurodevelopmental disorders that result in variable epilepsy, ataxia, spasticity, growth abnormalities, dysmorphism, hearing, and visual impairments, and a number of missense variants have been observed to be at intermediate allele frequencies (Yoon et al. 2017; Yap et al. 2021; Lin et al. 2023), a pattern potentially consistent with the action of balancing selection (as well as consistent with the observed CLR value).

Genomic segments of ≥50bp in length that alter genomic organization are known collectively as structural variants. Recent work by Versoza et al. (2026c) characterized the landscape of structural variation in the coppery titi monkey genomes, identifying over 13,000 such variants, including insertions, duplications, inversions, and deletions. We identified four of these deletions as strong balancing selection candidates based on our genome scan results (see Table 1 and Figure 5). All four deletions reside in genomic regions lacking functional annotation; moreover, three reside in regions of the genome with relatively reduced recombination rates (Soni, Versoza et al. 2026), though that observation may itself owe to the presence of the segregating variant. Thus, while this is an intriguing observation, insufficient evidence is available to further interpret these structural variant results.

**Table 1:** Genomic coordinates of deletions identified as balancing selection candidates. The maximum composite likelihood ratio (CLR) value is the highest CLR inferred by *B_0MAF_* for SNPs that are located within the deletion. Recombination rates are calculated as the mean rate across each deletion, from fine-scale recombination maps inferred by Soni, Versoza et al. (2026).

| chromosome | start | end | length | maximum<br>CLR | mean<br>recombination<br>rate (cM/Mb) |
| --- | --- | --- | --- | --- | --- |
| 2 | 23,269,995 | 23,273,368 | 3,373 | 240.28 | 0.426 |
| 9 | 21,882,705 | 21,885,776 | 3,071 | 300.37 | 0.035 |
| 14 | 58,818,610 | 58,819,850 | 1,240 | 329.20 | 0.467 |
| 20 | 16,496,560 | 16,497,174 | 614 | 296.54 | 8.087 |

## CONCLUSION

In this study, we have characterized long-term selective dynamics in the coppery titi monkey genome by fitting a DFE to observed patterns of polymorphism and divergence, finding evidence of an increased proportion of effectively neutral mutations relative to both humans and common marmosets. Additionally, applying the first genome-wide scans for positive and balancing selection in the coppery titi monkey genome based on an evolutionary baseline model incorporating the species-specific population history, mutation rates, and recombination rates, we identified a small number of promising candidate loci, several of which are of notable biomedical significance.

## MATERIALS AND METHODS

### Animal Subjects

Animals were maintained at the California National Primate Research Center (Davis, CA). This study was performed in compliance with all regulations regarding the care and use of captive primates, including the NIH Guidelines for the Care and Use of Animals and the American Society of Primatologists’ Guidelines for the Ethical Treatment of Nonhuman Primates. Procedures were approved by the UC-Davis Institutional Animal Care and Use Committee (protocol 22523).

### Whole genome, population-level data

We isolated genomic DNA from peripheral blood samples of six unrelated coppery titi monkey individuals, obtained during routine veterinary examinations. For each sample, we prepared a 150 bp paired-end sequencing library using a PCR-free protocol, and sequenced the resulting libraries on an Illumina NovaSeq 6000 system (Illumina, San Diego, CA, USA). We trimmed remnant adapter sequences from the 3’-end of the raw reads using fastp v.0.24.0 (Chen et al. 2018). Using BWA-MEM v.0.7.15 (Li 2013), we aligned the adapter-trimmed reads to the coppery titi monkey genome assembly (GenBank #: GCA_040437455.1; Pfeifer et al. 2024), marking split reads (*-M*). In line with established best-practice workflows (Pfeifer 2017), we first identified and flagged read duplicates using the *MarkDuplicates* module available via the Genome Analysis Toolkit (GATK) v.4.4.0 (van der Auwera and O’Connor 2020) to eliminate high-coverage support arising from optical and sampling biases, and then performed base quality score recalibration using the *BaseRecalibrator* and *ApplyBQSR* modules to correct for systematic biases in machine-reported base quality scores. Recalibrations were based on a high-confidence training dataset of graph-based, high-precision genotypes previously obtained from the colony (Versoza et al. 2026a). Using these pre-processed read mappings, we called both polymorphic and monomorphic sites in each sample using the *HaplotypeCaller* module (with ‘*-ERC* BP_RESOLUTION’ enabled), applying a minimum mapping quality threshold of 40 (’*--minimum-mapping-quality* 40’) and disabling PCR error correction (’*-pcr-indel-model* NONE’). Afterward, we combined per-sample outputs using the *CombineGVCFs* module and performed joint genotyping across all samples using the *GenotypeGVCFs* module (with the ‘*-all-sites*’ option enabled). To improve genotyping accuracy, polymorphic sites were re-genotyped by generating local variation graphs with Graphtyper v.2.7.2 (Eggertsson et al. 2017).

We focused our downstream analyses on autosomal, biallelic SNPs and monoallelic sites genotyped in every sample that passed built-in quality criteria and that were located outside of regions exhibiting an unusually depth of coverage (defined here as <0.5× or >2× the mean depth per sample), harboring nearby insertions or deletions (within 5 bp), or showing a low mappability (based on a 150bp-read SNPable mask with a stringency parameter of 1; https://lh3lh3.users.sourceforge.net/snpable.shtml).

### Calculating exonic divergence between coppery titi monkeys and humans

To calculate divergence between *P. cupreus* and *H. sapiens*, we replaced the scaffold-level *P. cupreus* genome assembly contained in the 447-way multiple species alignment (Zoonomia Consortium 2020) with the highly contiguous, chromosome-level, annotated genome assembly of Pfeifer et al. (2024), following the approach of Soni, Versoza et al. (2026) via HAL v.2.2 (Hickey et al. 2013) and Cactus v.2.9.2 (Armstrong et al. 2020). We then extracted the sub-alignment containing the updated coppery titi monkey genome, the human genome, and the reconstructed ancestor (PrimatesAnc003), identifying fixed differences between coppery titi monkeys and humans via HAL’s *halSnps* function. Specifically, we focused on those fixed differences where coppery titi monkeys exhibited a different allele than humans and PrimatesAnc003, with the latter two sharing the same allele. We excluded sites containing gaps or missing nucleotides, and limited our analyses to regions in which we could confidently map the short-read data (see the “Whole genome, population-level data” section for details). These cautionary steps are necessary due to the relatively long divergence time between humans and coppery titi monkeys. Finally, we extracted exonic sites in order to calculate divergence per exon.

### DFE inference

We ran forward-in-time simulations in SLiM v.5.0.1 (Haller et al. 2026), under the demographic model of Terbot et al. (2026) inferred using a mutation rate of 1.07 × 10^-8^ per base pair per generation (/bp/gen) (referred to as the high mutation rate model throughout this study). Briefly, an ancestral population of size *N_ancestral_* = 17,807 individuals underwent a population contraction to ∼35,000 individuals 122,126 generations ago, before expanding to ∼425,000 individuals 55,881 generations ago. Finally, the population underwent a severe contraction 1,406 generations ago, to the current population size of 6,330 individuals. Following a 10 *N_ancestral_* burn-in, we simulated a divergence phase between *P. cupreus* and humans of 32 million years (Glazko and Nei 2003) — assuming a generation time of 6 years (Pacifici et al. 2013; Perez et al. 2013) — after which the demographic model was simulated. In brief, 100 replicates of a 618bp region were simulated; this region length is the mean empirical exon length, when limiting to those exons > 500bp in length. We drew mutation and recombination rates from a normal distribution for each simulated exon, using a mean mutation rate of 1.07 × 10^-8^ /bp/gen and a mean recombination rate of 1.00 × 10^-8^ /bp/gen (i.e., the rates used by Terbot et al. [2026] to infer the *P. cupreus* demographic model), such that the mean rate across all 100 simulation replicates was equal to these mean rates. We drew mutations from a DFE comprised of three fixed classes (Johri et al. 2020): nearly neutral mutations (2*N*_ancestral_ *s* < 10, where *N_ancestral_* is the ancestral population size and *s* is the reduction in fitness of the mutant homozygote relative to the wild-type), weakly/moderately deleterious mutations (10 ≤ 2*N*_ancestral_ *s* < 100), and strongly deleterious mutations (100 ≤ 2*N*_ancestral_ *s*).

To infer the best-fitting DFE, we performed a grid search by simulating under varying proportions of mutations in each DFE category and comparing the fit of the mean and variance of four summary statistics between our empirical and 100 simulation replicates: Watterson’s θ_w_ (Watterson 1975) per site, Tajima’s *D* (Tajima 1989) per 10kb, the number of singletons per site, and exonic divergence per site. We used pylibseq v.1.8.3 (Thornton 2003) to calculate all summary statistics except exonic divergence, which was calculated as the number of fixations that occurred in our simulated population post-burn-in. This approach enabled direct comparison to the empirically observed number of substitutions along the *P. cupreus* branch, following the split from humans. Starting with the DFEs inferred by Johri et al. (2023) for humans and Soni et al. (2025c) for the common marmoset (i.e., DFEs inferred using a similar approach in a species with a similar effective population size as well as another platyrrhine, respectively), we performed manual grid step iterations based upon the fit of the DFE to the data, until a visually consistent fit of the mean and variance of our chosen summary statistics between our empirical and simulated data was identified.

### Power analyses

To evaluate how much statistical power exists to detect episodic selective events in our *P. cupreus* population, we performed 100 simulation replicates under the high mutation rate demographic model inferred by Terbot et al. (2026; see the “DFE inference” section for details), using a 10-fold scaling reduction in population size. We simulated 33 functional regions — comprised of nine 130bp exons, separated by introns of size 1591bp. Functional regions were flanked by intergenic regions of length 16,489bp (following Soni et al. 2025b). The number of functional regions chosen were such that the total simulated length was ∼1Mb. Exonic mutations were drawn from the DFE inferred in this study, whilst intronic and intergenic mutations were selectively neutral. In order to model mutation rate heterogeneity, mutation rates were drawn from the empirical fine-scale mutation rate distribution inferred by Soni, Versoza et al. (2026) for each 1kb window, such that the mean rate across each simulated replicate was equal to the (10-fold rescaled) mean rate of 1.07 × 10^-7^ /bp/gen. Recombination rates were drawn from a normal distribution with a mean of 1.00 × 10^-7^ /bp/gen, the mean rescaled rate across all windows per simulation replicate. In each replicate, a single positively selected mutation was introduced, with two possible beneficial population-scaled selection coefficients 2*N_ancestral_ s* = [100, 1000], and four times of introduction of the beneficial mutation in *N_ancestral_* generations *1* = [0.1, 0.2, 0.5, 1.0], for selective sweep analyses. For balancing selection analysis, we introduced the balanced mutation at *1_b_* = [10, 50, 75] *N_ancestral_* generations prior to the end of the simulation (i.e., representing time segregating prior to sampling). The balanced mutation experienced negative-frequency dependent selection, with the selection coefficient of the balanced mutation, *S_bp_*, dependent on its frequency in the population (*F_eq_*) and its equilibrium frequency (here modelled as 0.5): *S_bp_* = 0.5 – *F_eq_*. Simulations were restarted if the beneficial mutation failed to fix or was lost from the population in the case of selective sweeps, or if the balanced mutation was fixed or lost from the population (restarting at the time of introduction of the balanced mutation).

To quantify statistical power, we performed genome scans on this simulated data using SweepFinder2 (DeGiorgio et al. 2016) and *B_0MAF_* (Cheng and DeGiorgio 2020). SweepFinder2 inference was performed at each SNP, whilst *B_0MAF_* inference was performed at every fifth SNP. The following SweepFinder2 command was used: *SweepFinder2 –lu GridFile FreqFile SpectFile OutFile*. For *B_0MAF_* the following command was used: *python BalLerMix+_v1.py –i InputFile –-spect SfsFile –o OutFile –s 5 –-usePhysPos –MAF –-noSub –-rec 1e-5* ROC plots were generated by combining loci into genomic windows of sizes [100, 1,000, 10,000, 100,000] bp for selective sweep inference, and sizes [1,000, 10,000, 100,000] bp for balancing selection inference, with the highest CLR per window utilized for assessing whether the window was a true positive, false positive, true negative, or false negative.

### Generating null thresholds for genome scans of recent positive selection and balancing selection

We simulated the two *P. cupreus* demographic models inferred by Terbot et al. (2026) in order to generate genome-wide null thresholds for selection inference. For details of the demographic model inferred using the higher mutation rate of 1.07 × 10^-8^ /bp/gen, see the section above entitled “DFE inference”. The demographic model inferred using the lower end of mutation rate estimates (0.488 × 10^-8^ /bp/gen) — generated for comparison and to account for underlying uncertainty — also involves three instantaneous population size changes. However, the ancestral population size in this model is *N_ancestral_* = 169,079, with a population contraction to ∼45,000 individuals occurring 351,168 generations ago, followed by an expansion to almost 2 million individuals that occurred 131,015 generations ago. Finally, under this model, the population undergoes a severe population contraction 3,160 generations ago, to the present-day size of 12,338 individuals. These demographic models were inferred on a dataset in which functional regions, as well as the 10kb flanking functional regions, were masked to avoid the biasing effects of purifying selection and BGS (Charlesworth et al. 1993; Johri et al. 2020). We therefore simulated the full masked length of each chromosome, with 10 replicates simulated for each chromosome. We drew mutation and recombination rates from a uniform distribution for each simulated replicate, using a mean mutation rate of 1.07 × 10^-8^ /bp/gen or 0.488 × 10^-8^ /bp/generation for the respective demographic model, and a mean recombination rate of 1.00 x 10^-8^ /bp/gen, such that the mean rate per simulation replicate was equal to these mean rates.

To generate genome-wide null thresholds, we ran SweepFinder2 (DeGiorgio et al. 2016) and the *B_0MAF_* balancing selection approach (Cheng and Degiorgio 2020) on our simulated data. We ran each software using the same commands outlined in the preceding section (see the section entitled “Power analyses”). Because we lacked information on the polarization of SNPs, allele frequencies were folded, and only polymorphic sites were considered. The highest CLR value across all null model simulations was set as the null threshold for selection inference, given that this observation represents the highest value generated in the absence of selective sweeps or balancing selection, thereby minimizing the downstream number of false positives.

### Inferring recent positive selection and balancing selection in the *P. cupreus* genome

We applied the inference approach discussed in the preceding section on our empirical data, utilizing the same commands used for the generation of null thresholds and for power analyses. Loci at which the empirical CLR was greater than the null threshold value were considered putative selection candidates. We identified candidate genes in which these loci were located. To verify the quality of the gene annotations for these candidate genes, we ran a BLASTx (Altschul et al. 1990) search on the coding sequences of our candidate genes to identify the best hits in humans. We then ran BLASTp on the best hits to verify whether the best hit in coppery titi monkeys returned the original sequence. We also compared the number of conserved domains across all protein coding sequences in a given gene, to the number of conserved domains in the best hits in humans, using CD-Search (Marchler-Bauer and Bryant 2004). Finally, we checked for synteny by comparing the flanking genes for each candidate gene between coppery titi monkeys and humans. Only genes with an amino acid identity >85%, sequence divergence <15%, close alignment between the number of conserved domains between coppery titi monkeys and humans, and synteny were considered for functional analysis.

We manually curated gene functions and expressions via the NCBI database (Sayers et al. 2022) and the Expression Atlas (Madeira et al. 2022) respectively. Finally, we performed gene ontology analysis (The Gene Ontology Consortium 2023) via the Database for Annotation, Visualization, and Integrated Discovery (DAVID; Ma et al. 2023).

## ACKNOWLEDGEMENTS

We would like to thank the California National Primate Research Center for providing the coppery titi monkey samples used in this study. Computations were performed on the Open Science Grid (OSG 2015) supported by National Science Foundation awards 2030508 and 2323298, the JetStream2 supercomputer at Indiana University through allocation BIO250421 from the ACCESS program (Boerner et al. 2023) supported by National Science Foundation awards 2138259, 2138286, 2138307, 2137603, and 2138296, and the Sol supercomputer at Arizona State University (Jennewein et al. 2023).

## FUNDING

This work was supported by the National Institute of General Medical Sciences of the National Institutes of Health under Award Number R35GM151008 to SPP, Award Number R35GM139383 to JDJ, and the California National Primate Research Center base grant P51OD011107. CJV was supported by the National Science Foundation CAREER Award DEB-2045343 to SPP. KLB was supported by the Eunice Kennedy Shriver National Institute of Child Health and Human Development and the National Institute of Mental Health of the National Institutes of Health under Award Numbers R01HD092055 and MH125411, and by the Good Nature Institute. The content is solely the responsibility of the authors and does not necessarily represent the official views of the National Institutes of Health or the National Science Foundation.

## Notes

### Competing Interest Statement

The authors have declared no competing interest.

