## Supplementary Materials for "Inferring patterns of purifying, positive and balancing selection in the coppery titi monkey (*Plecturocebus cupreus*) utilizing a well-fit evolutionary baseline model"

**Table S1**

| GO Term | Genes | Count | List Total | P-Value | Benjamini | Fold Enrichment | Bonferroni | FDR | Fisher Exact |
|---|---|---|---|---|---|---|---|---|---|
| acetyltransferase activator activity | 10.53% | 2 | 19 | 1.02E-02 | 6.14E-01 | 184.42 | 4.60E-01 | 6.14E-01 | 5.04E-05 |
| cytosol | 52.63% | 10 | 19 | 3.36E-02 | 1.00E+00 | 1.93 | 8.88E-01 | 1.00E+00 | 1.68E-02 |
| receptor tyrosine kinase binding | 10.53% | 2 | 19 | 6.08E-02 | 1.00E+00 | 30.28 | 9.77E-01 | 1.00E+00 | 1.96E-03 |
| insulin receptor signaling pathway | 10.53% | 2 | 19 | 7.13E-02 | 1.00E+00 | 25.67 | 1.00E+00 | 1.00E+00 | 2.71E-03 |
| cytoplasm | 57.89% | 11 | 19 | 8.81E-02 | 1.00E+00 | 1.55 | 9.97E-01 | 1.00E+00 | 5.42E-02 |

**Table S1: Results of gene functional analysis using DAVID (Ma et al. 2023) for the 19 selective sweep candidate genes.**

**Table S2**

| Term | Genes | Count | List Total | P-Value | Benjamini | Fold Enrichment | Bonferroni | FDR | Fisher Exact |
|---|---|---|---|---|---|---|---|---|---|
| C2_domain_sf | 17.65% | 3 | 15 | 6.11E-3 | 4.89E-1 | 23.53 | 3.88E-1 | 4.89E-1 | 2.55E-4 |
| Host-virus interaction | 23.53% | 4 | 9 | 9.58E-3 | 1.05E-1 | 7.41 | 1.00E-1 | 1.05E-1 | 1.27E-3 |
| Repeat | 52.94% | 9 | 12 | 1.24E-2 | 1.11E-1 | 2.14 | 1.06E-1 | 1.11E-1 | 5.68E-3 |
| nucleoplasm | 35.29% | 6 | 13 | 6.81E-2 | 1.00E+0 | 2.35 | 9.86E-1 | 1.00E+0 | 2.77E-2 |
| nucleus | 47.06% | 8 | 13 | 7.22E-2 | 1.00E+0 | 1.82 | 9.90E-1 | 1.00E+0 | 3.83E-2 |
| Ubiquitin mediated proteolysis | 11.76% | 2 | 6 | 7.26E-2 | 1.00E+0 | 22.29 | 8.59E-1 | 1.00E+0 | 3.20E-3 |
| Tight junction | 11.76% | 2 | 14 | 7.65E-2 | 5.44E-1 | 23.42 | 7.41E-1 | 5.44E-1 | 3.20E-3 |
| bicellular tight junction | 11.76% | 2 | 13 | 7.68E-2 | 1.00E+0 | 23.2 | 9.92E-1 | 1.00E+0 | 3.25E-3 |
| endosome | 17.65% | 3 | 13 | 7.68E-2 | 1.00E+0 | 5.94 | 9.92E-1 | 1.00E+0 | 1.25E-2 |
| Endosome | 17.65% | 3 | 14 | 8.88E-2 | 5.44E-1 | 5.51 | 7.94E-1 | 5.44E-1 | 1.55E-2 |
| C2 | 11.76% | 2 | 9 | 9.04E-2 | 1.00E+0 | 18.88 | 8.00E-1 | 1.00E+0 | 4.69E-3 |
| DOMAIN:C2 | 11.76% | 2 | 16 | 9.43E-2 | 1.00E+0 | 19 | 1.00E+0 | 1.00E+0 | 4.85E-3 |
| C2_dom | 11.76% | 2 | 15 | 9.50E-2 | 1.00E+0 | 18.76 | 1.00E+0 | 1.00E+0 | 4.96E-3 |
| Nucleus | 47.06% | 8 | 14 | 9.60E-2 | 5.44E-1 | 1.74 | 8.20E-1 | 5.44E-1 | 5.26E-2 |

**Table S2: Results of gene functional analysis using DAVID (Ma et al. 2023) for the 19 most rapidly diverging genes.** These are genes whose mean divergence across exons is greater than the 99.9$^{th}$ percentile neutral divergence between coppery titi monkeys and humans, as inferred by Soni, Versoza et al. (2026).

**Supplementary Figure S1**

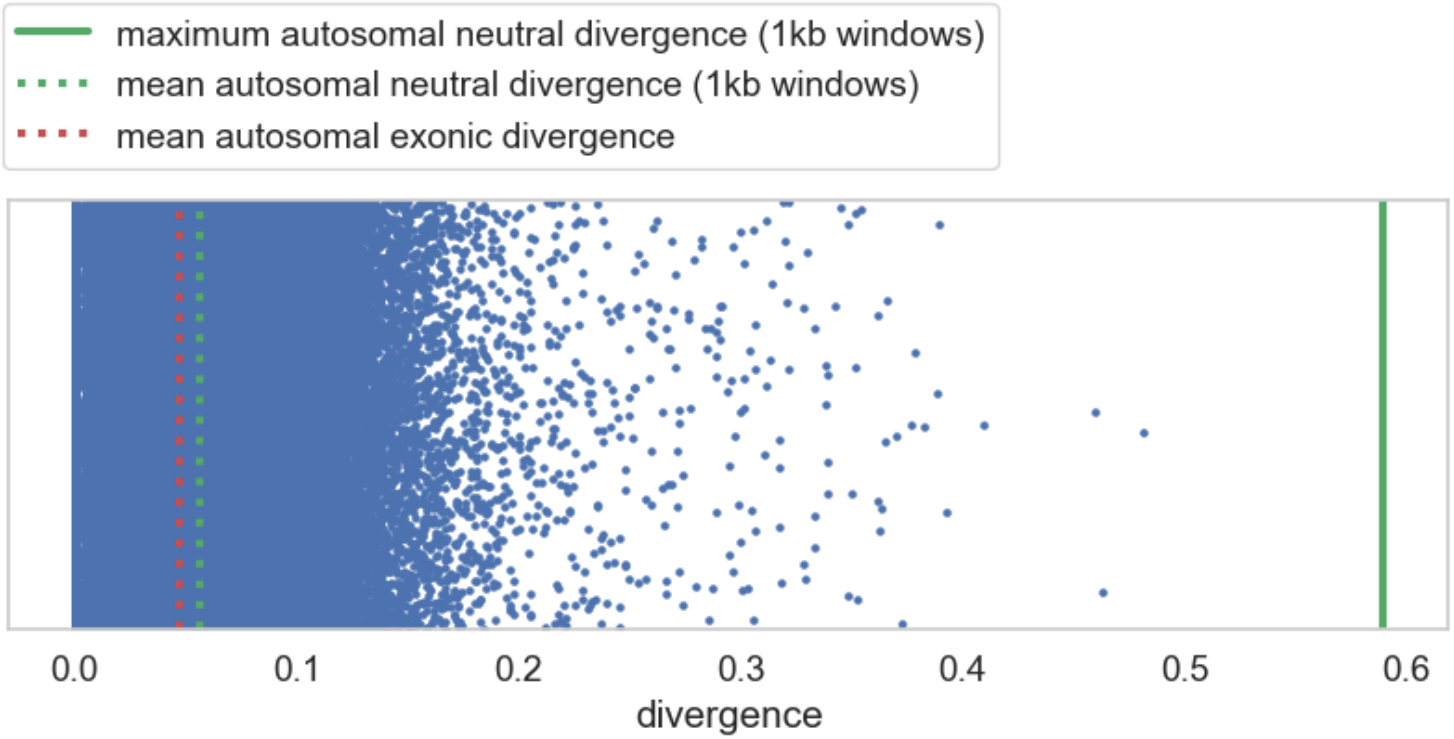


**Supplementary Figure S1:** Exonic divergence between the coppery titi monkey and humans. Scatter plot with maximum neutral divergence values marked for windows of size 1 kb (green solid line), as well as the mean neutral divergence for 1 kb windows (green dashed line), as calculated from non-functional autosomal regions of the coppery titi monkey genome (see Soni, Versoza et al. 2026). Each dot represents an autosomal exon, and the mean exonic divergence as calculated in this study is plotted (red dashed line).

**Supplementary Figure S2**

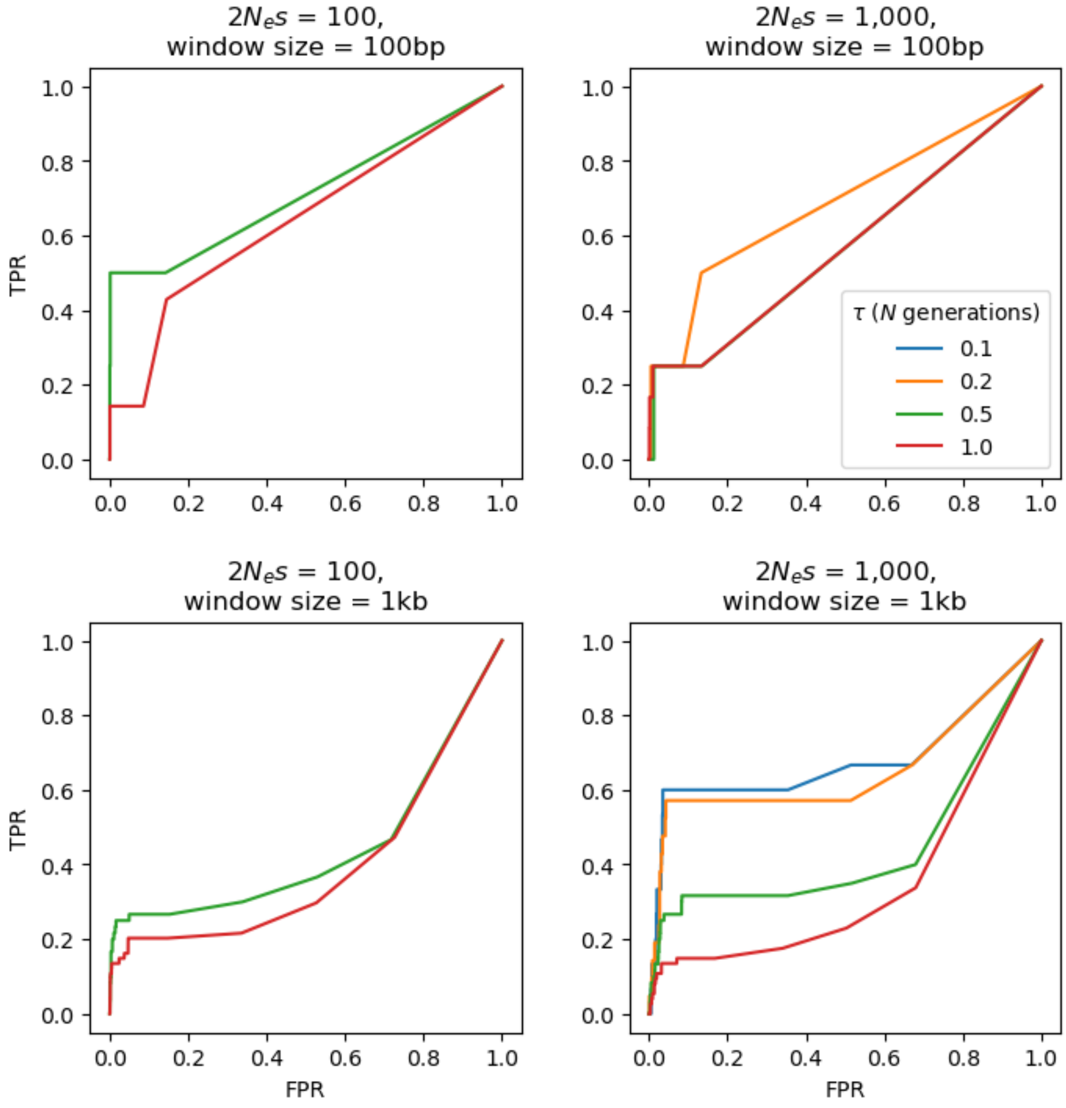


**Supplementary Figure S2: ROC plots based on 100 simulated replicates under the coppery titi monkey demographic model inferred by Terbot et al. (2026) for selective sweep inference using the SweepFinder2 method.** The false-positive rate (FPR) and the true-positive rate (TPR) are provided on the *x*-axis and *y*-axis of the ROC plots, respectively. Power analyses were conducted across two selection regimes — population-scaled strengths of selection of $2N_es$ = 100 and 1000; four times of introduction of the beneficial mutation ($\tau$ = 0.1, 0.2, 0.5, and 1.0 *N* generations ago), and window sizes of 100bp and 1kb. Note that no ROC plots could be plotted for simulation regimes in which no beneficial mutations reached fixation at the time of sampling.

**Supplementary Figure S3**

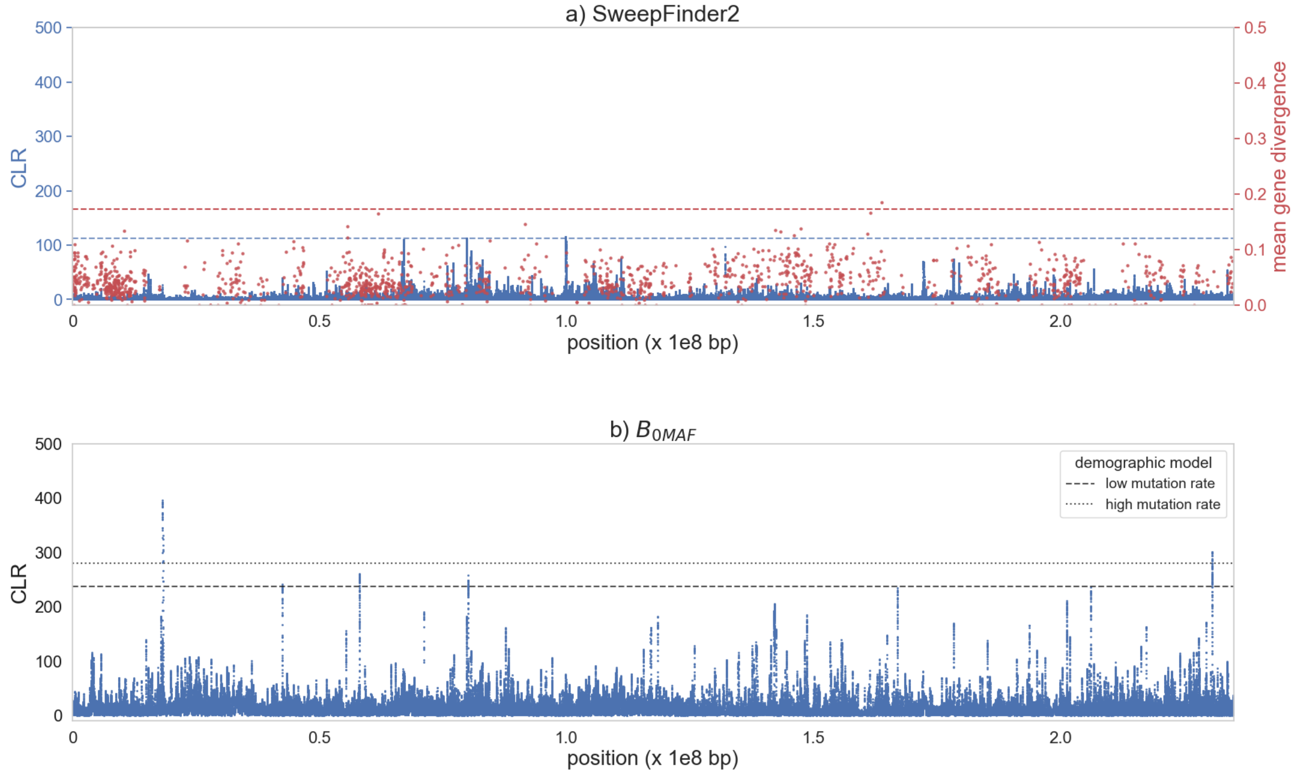


**Supplementary Figure S3: Genome scan results for chromosome 1 using a) SweepFinder2 (shown in blue) and empirical exonic divergence (red); and b) $B_{0MAF}$. a)** The x-axis shows the position along the chromosome, the left y-axis shows the composite likelihood ratio (CLR) value of the sweep statistic at each SNP, and the right y-axis provides the mean gene divergence. The horizontal blue dashed line represents the null threshold for sweep detection, and the horizontal red dashed line represents the 99.9$^{th}$ percentile neutral divergence. **b)** The x-axis shows the position along the chromosome, and the y-axis the CLR value of the balancing selection statistic at each fifth SNP. The horizontal dashed lines represent the null thresholds for detection of balancing selection, based on two different coppery titi demographic models (see the “Materials and Methods” section for details).

**Supplementary Figure S4**

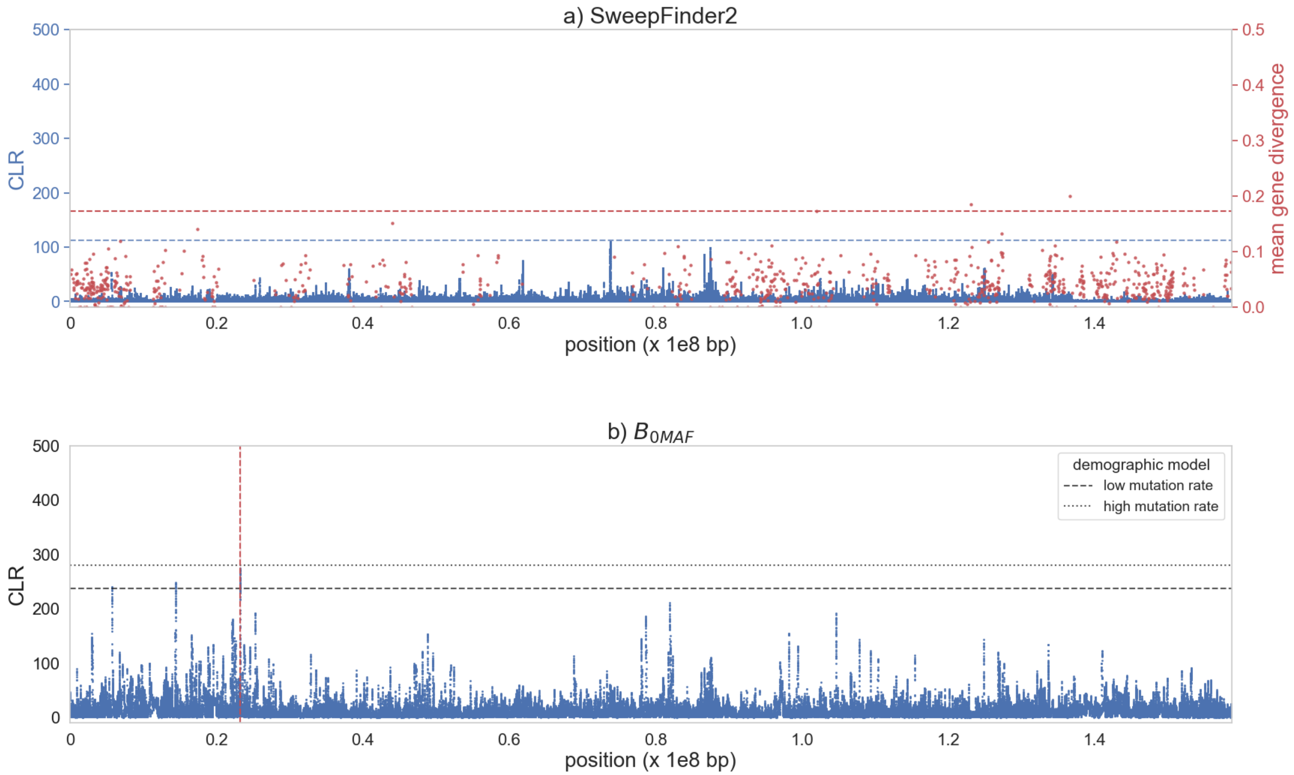


**Supplementary Figure S4: Genome scan results for chromosome 2 using a) SweepFinder2 (shown in blue) and empirical exonic divergence (red); and b) $B_{0MAF}$. a)** The x-axis shows the position along the chromosome, the left y-axis shows the composite likelihood ratio (CLR) value of the sweep statistic at each SNP, and the right y-axis provides the mean gene divergence. The horizontal blue dashed line represents the null threshold for sweep detection, and the horizontal red dashed line represents the 99.9th percentile neutral divergence. **b)** The x-axis shows the position along the chromosome, and the y-axis the CLR value of the balancing selection statistic at each fifth SNP. The horizontal dashed lines represent the null thresholds for detection of balancing selection, based on two different coppery titi demographic models (see the "Materials and Methods" section for details). The position of the deletion is marked by the vertical dashed red line.

### Supplementary Figure S5

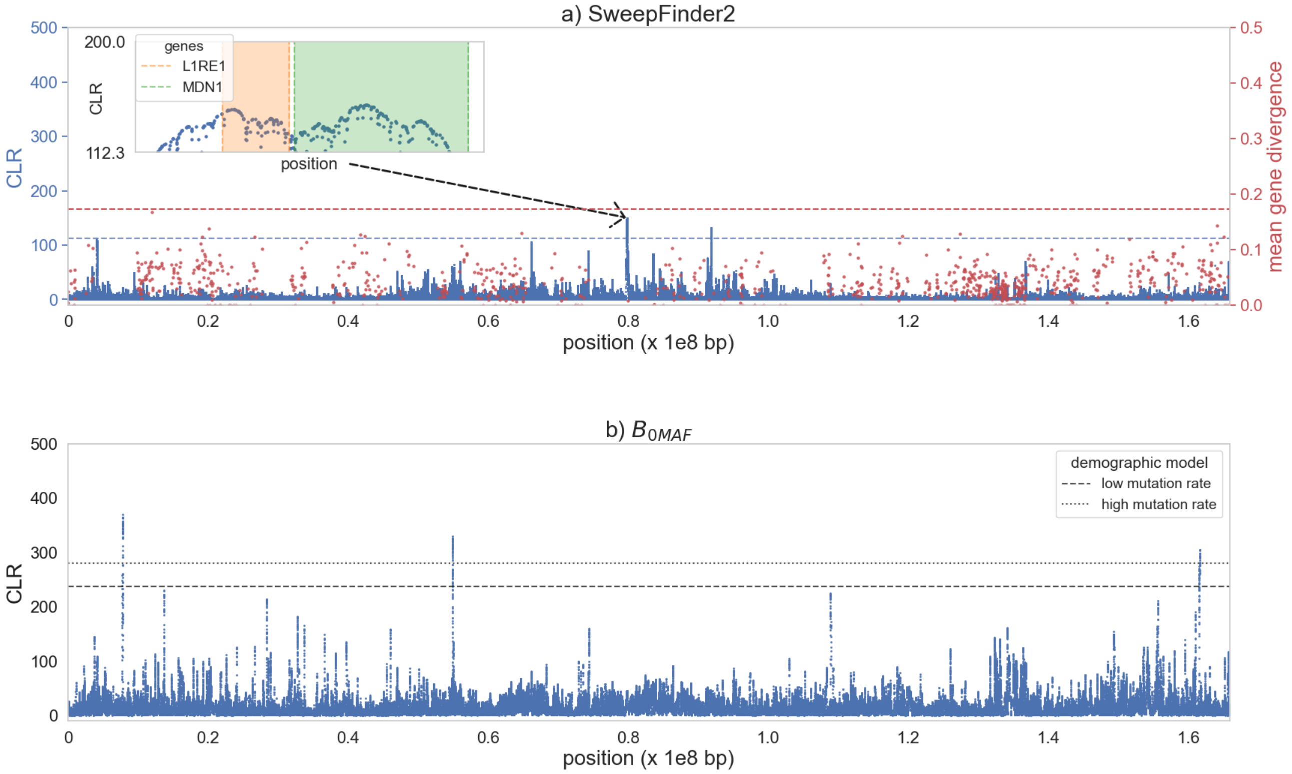


**Supplementary Figure S5: Genome scan results for chromosome 3 using a) SweepFinder2 (shown in blue) and empirical exonic divergence (red); and b) $B_{0MAF}$. a)** The x-axis shows the position along the chromosome, the left y-axis shows the composite likelihood ratio (CLR) value of the sweep statistic at each SNP, and the right y-axis provides the mean gene divergence. The horizontal blue dashed line represents the null threshold for sweep detection, and the horizontal red dashed line represents the 99.9$^{th}$ percentile neutral divergence. Inset plots zoom in on likelihood surface peaks, with genes in these regions highlighted. **b)** The x-axis shows the position along the chromosome, and the y-axis the CLR value of the balancing selection statistic at each fifth SNP. The horizontal dashed lines represent the null thresholds for detection of balancing selection, based on two different coppery titi demographic models (see the "Materials and Methods" section for details).

**Supplementary Figure S6**

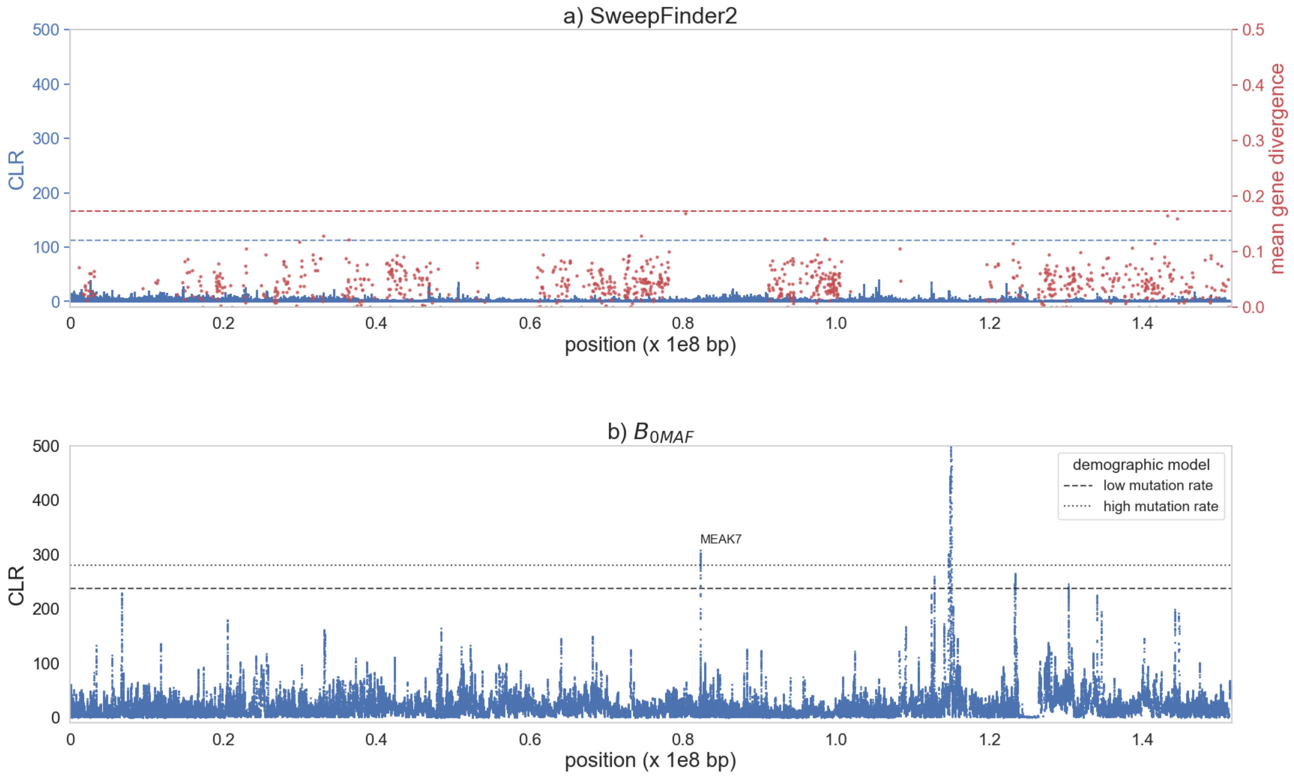


**Supplementary Figure S6: Genome scan results for chromosome 4 using a) SweepFinder2 (shown in blue) and empirical exonic divergence (red); and b) $B_{0MAF}$. a)** The x-axis shows the position along the chromosome, the left y-axis shows the composite likelihood ratio (CLR) value of the sweep statistic at each SNP, and the right y-axis provides the mean gene divergence. The horizontal blue dashed line represents the null threshold for sweep detection, and the horizontal red dashed line represents the 99.9$^{th}$ percentile neutral divergence. **b)** The x-axis shows the position along the chromosome, and the y-axis the CLR value of the balancing selection statistic at each fifth SNP. The horizontal dashed lines represent the null thresholds for detection of balancing selection, based on two different coppery titi demographic models (see the "Materials and Methods" section for details). Candidate genes are labeled.

**Supplementary Figure S7**

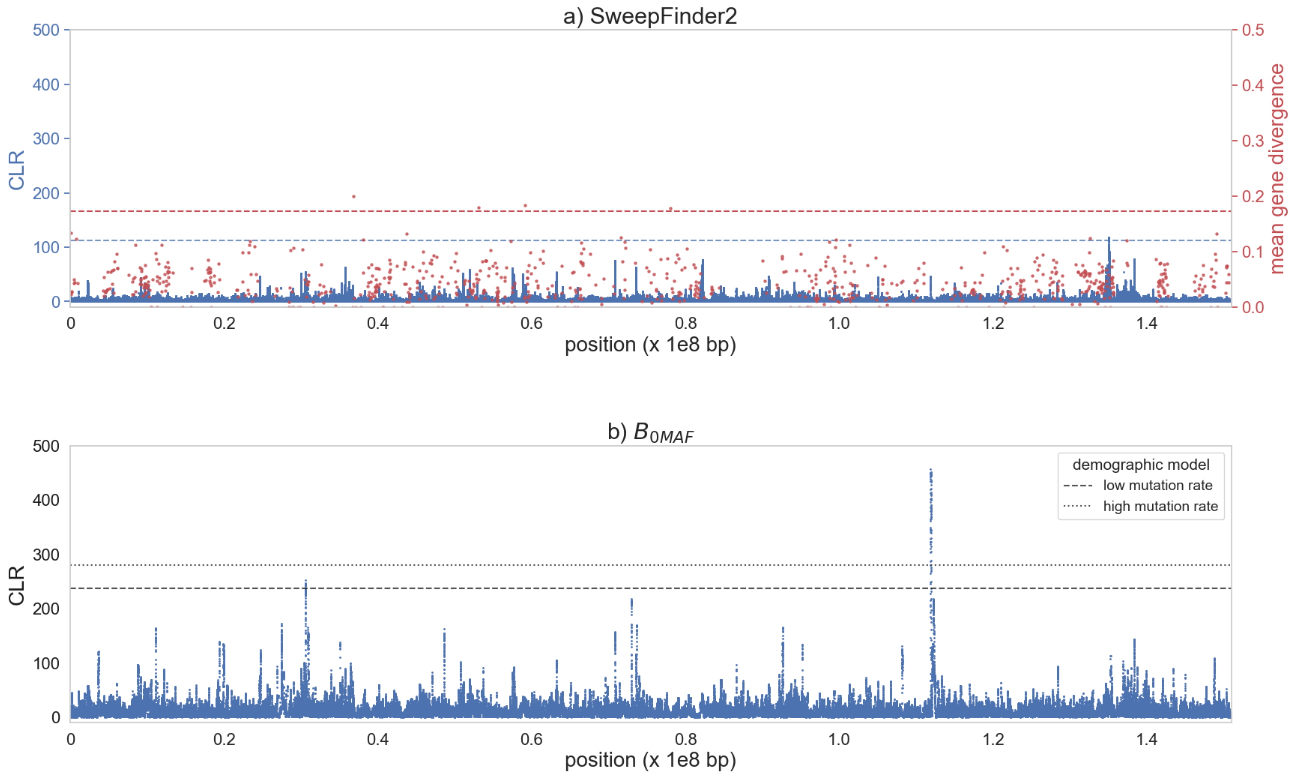


**Supplementary Figure S7: Genome scan results for chromosome 5 using a) SweepFinder2 (shown in blue) and empirical exonic divergence (red); and b) $B_{0MAF}$. a)** The x-axis shows the position along the chromosome, the left y-axis shows the composite likelihood ratio (CLR) value of the sweep statistic at each SNP, and the right y-axis provides the mean gene divergence. The horizontal blue dashed line represents the null threshold for sweep detection, and the horizontal red dashed line represents the 99.9$^{th}$ percentile neutral divergence. **b)** The x-axis shows the position along the chromosome, and the y-axis the CLR value of the balancing selection statistic at each fifth SNP. The horizontal dashed lines represent the null thresholds for detection of balancing selection, based on two different coppery titi demographic models (see the "Materials and Methods" section for details).

### Supplementary Figure S8

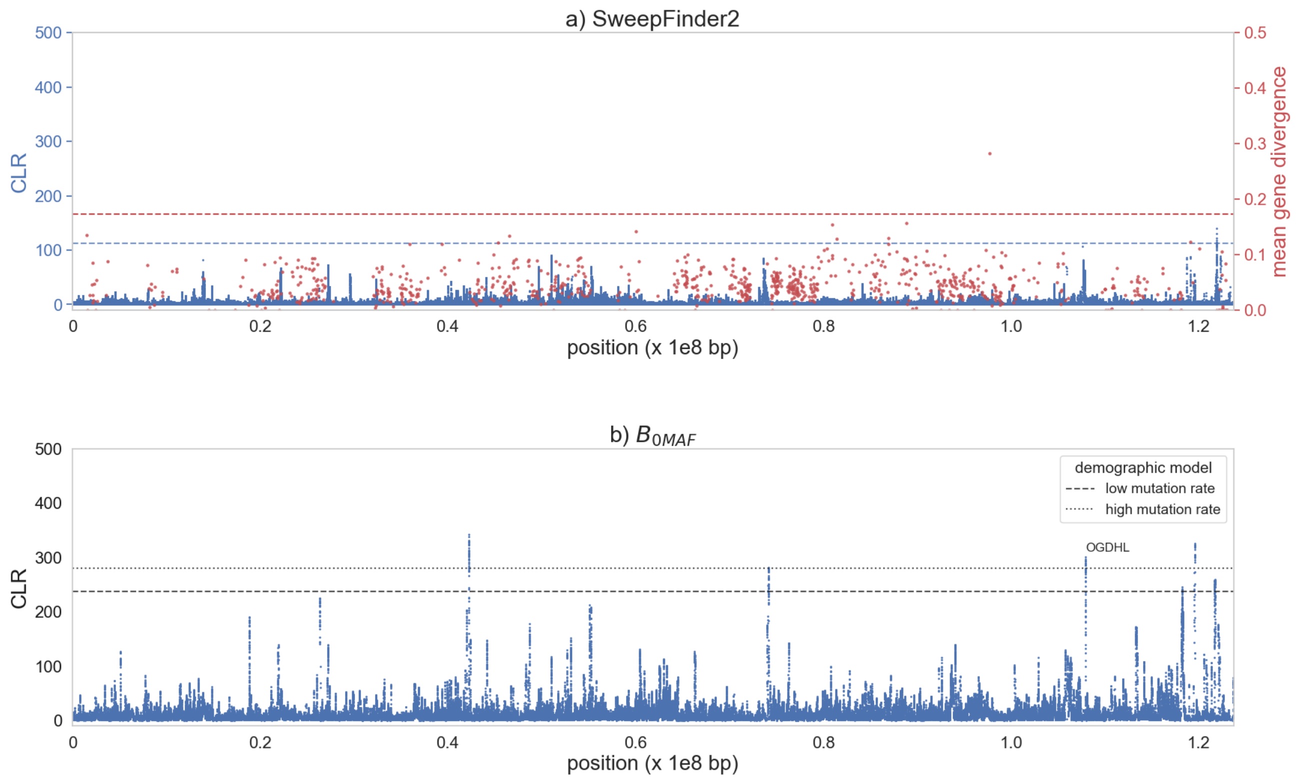


**Supplementary Figure S8: Genome scan results for chromosome 6 using a) SweepFinder2 (shown in blue) and empirical exonic divergence (red); and b) $B_{0MAF}$. a)** The x-axis shows the position along the chromosome, the left y-axis shows the composite likelihood ratio (CLR) value of the sweep statistic at each SNP, and the right y-axis provides the mean gene divergence. The horizontal blue dashed line represents the null threshold for sweep detection, and the horizontal red dashed line represents the 99.9$^{th}$ percentile neutral divergence. **b)** The x-axis shows the position along the chromosome, and the y-axis the CLR value of the balancing selection statistic at each fifth SNP. The horizontal dashed lines represent the null thresholds for detection of balancing selection, based on two different coppery titi demographic models (see the “Materials and Methods” section for details). Candidate genes are labeled.

**Supplementary Figure S9**

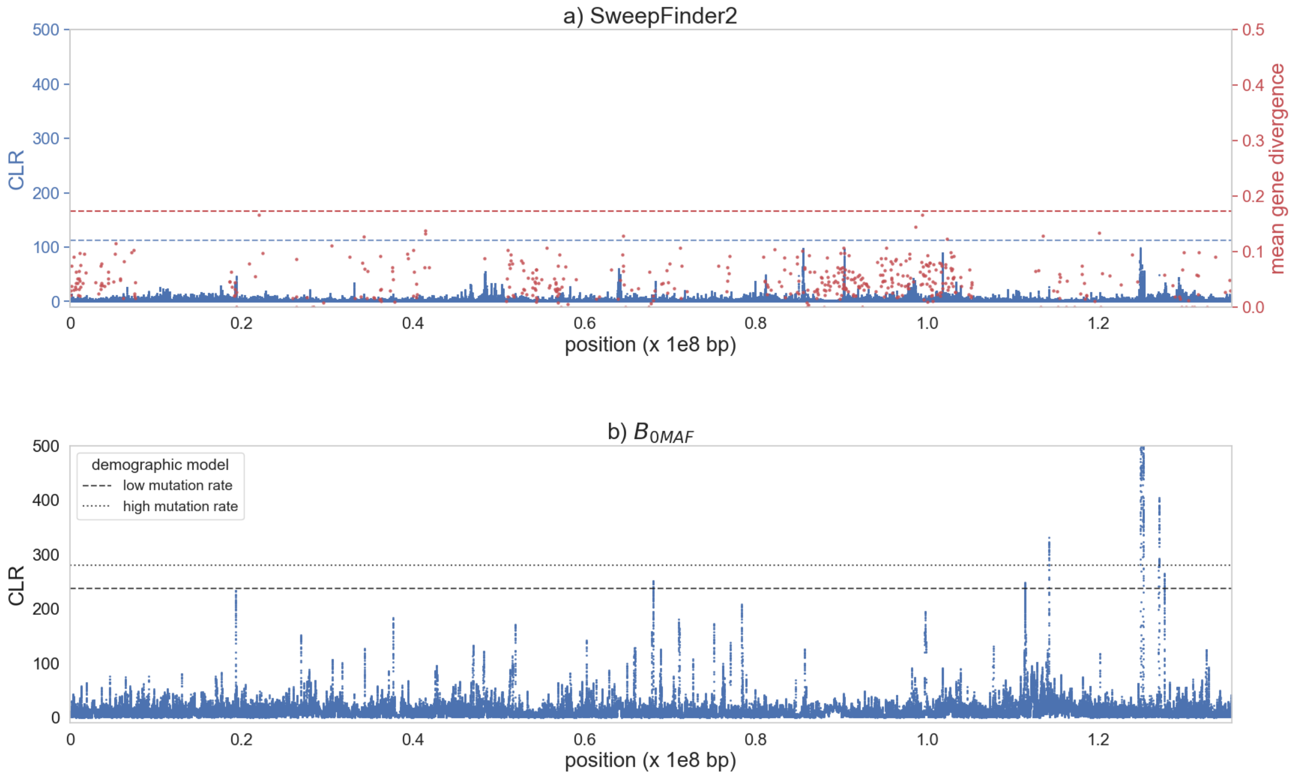


**Supplementary Figure S9: Genome scan results for chromosome 7 using a) SweepFinder2 (shown in blue) and empirical exonic divergence (red); and b) $B_{0MAF}$. a)** The x-axis shows the position along the chromosome, the left y-axis shows the composite likelihood ratio (CLR) value of the sweep statistic at each SNP, and the right y-axis provides the mean gene divergence. The horizontal blue dashed line represents the null threshold for sweep detection, and the horizontal red dashed line represents the 99.9$^{th}$ percentile neutral divergence. **b)** The x-axis shows the position along the chromosome, and the y-axis the CLR value of the balancing selection statistic at each fifth SNP. The horizontal dashed lines represent the null thresholds for detection of balancing selection, based on two different coppery titi demographic models (see the “Materials and Methods” section for details).

### Supplementary Figure S10

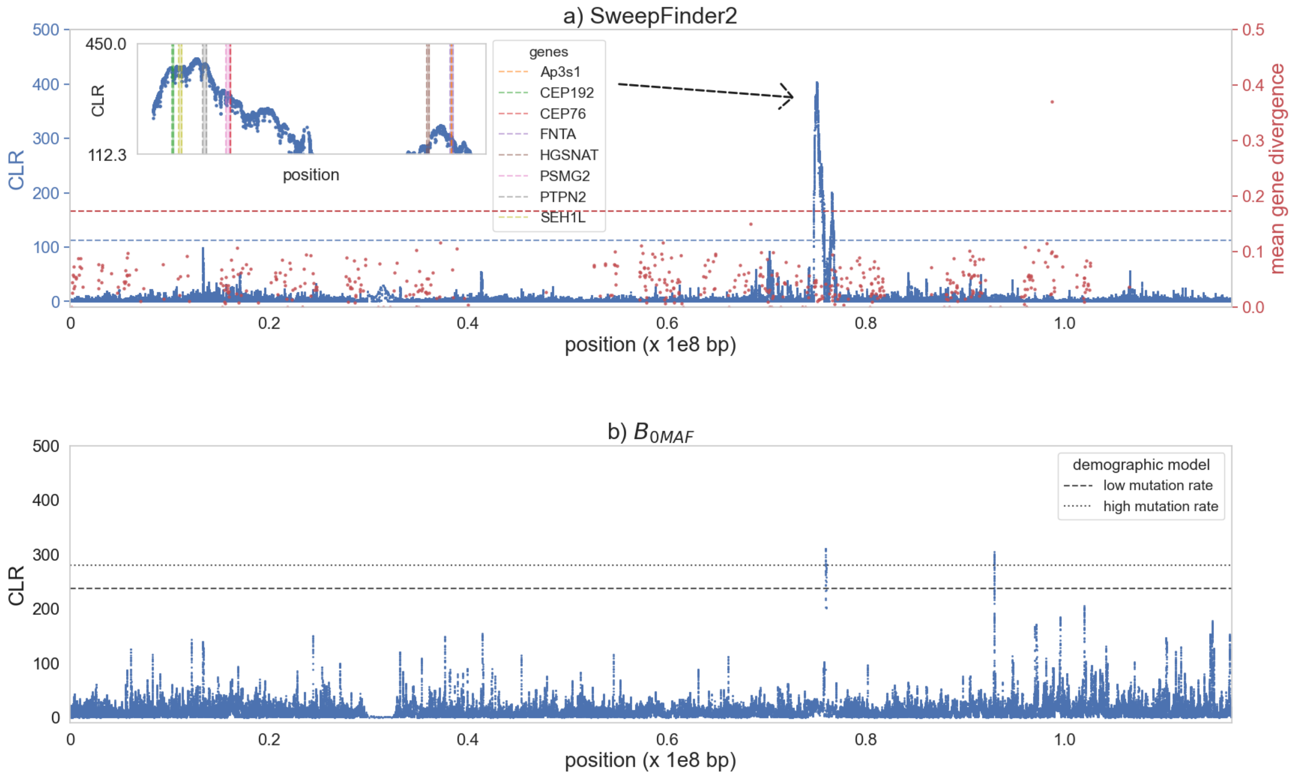


**Supplementary Figure S10: Genome scan results for chromosome 8 using a) SweepFinder2 (shown in blue) and empirical exonic divergence (red); and b) $B_{0MAF}$. a)** The x-axis shows the position along the chromosome, the left y-axis shows the composite likelihood ratio (CLR) value of the sweep statistic at each SNP, and the right y-axis provides the mean gene divergence. The horizontal blue dashed line represents the null threshold for sweep detection, and the horizontal red dashed line represents the 99.9th percentile neutral divergence. Inset plots zoom in on likelihood surface peaks, with genes in these regions highlighted. **b)** The x-axis shows the position along the chromosome, and the y-axis the CLR value of the balancing selection statistic at each fifth SNP. The horizontal dashed lines represent the null thresholds for detection of balancing selection, based on two different coppery titi demographic models (see the "Materials and Methods" section for details).

**Supplementary Figure S11**

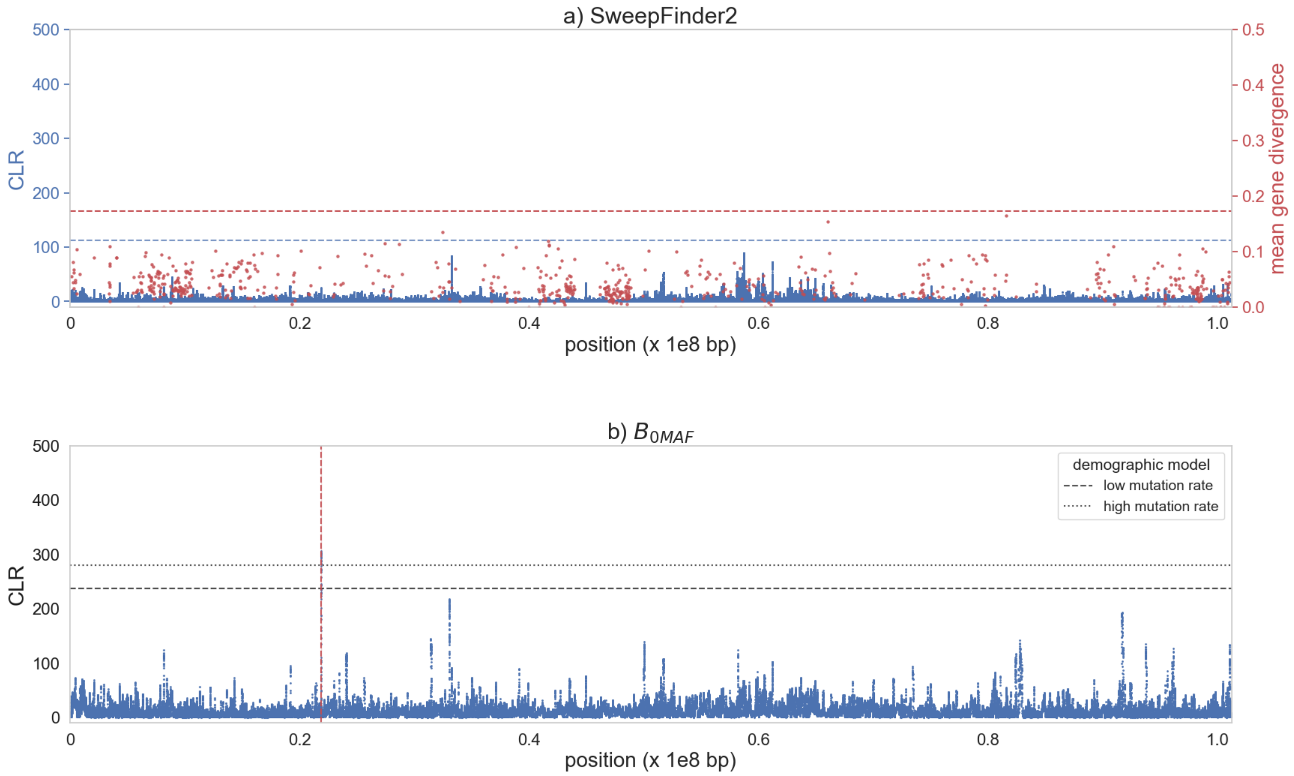


**Supplementary Figure S11: Genome scan results for chromosome 9 using a) SweepFinder2 (shown in blue) and empirical exonic divergence (red); and b) $B_{0MAF}$. a)** The x-axis shows the position along the chromosome, the left y-axis shows the composite likelihood ratio (CLR) value of the sweep statistic at each SNP, and the right y-axis provides the mean gene divergence. The horizontal blue dashed line represents the null threshold for sweep detection, and the horizontal red dashed line represents the 99.9$^{th}$ percentile neutral divergence. **b)** The x-axis shows the position along the chromosome, and the y-axis the CLR value of the balancing selection statistic at each fifth SNP. The horizontal dashed lines represent the null thresholds for detection of balancing selection, based on two different coppery titi demographic models (see the "Materials and Methods" section for details). The position of the deletion is marked by the vertical dashed red line.

**Supplementary Figure S12**

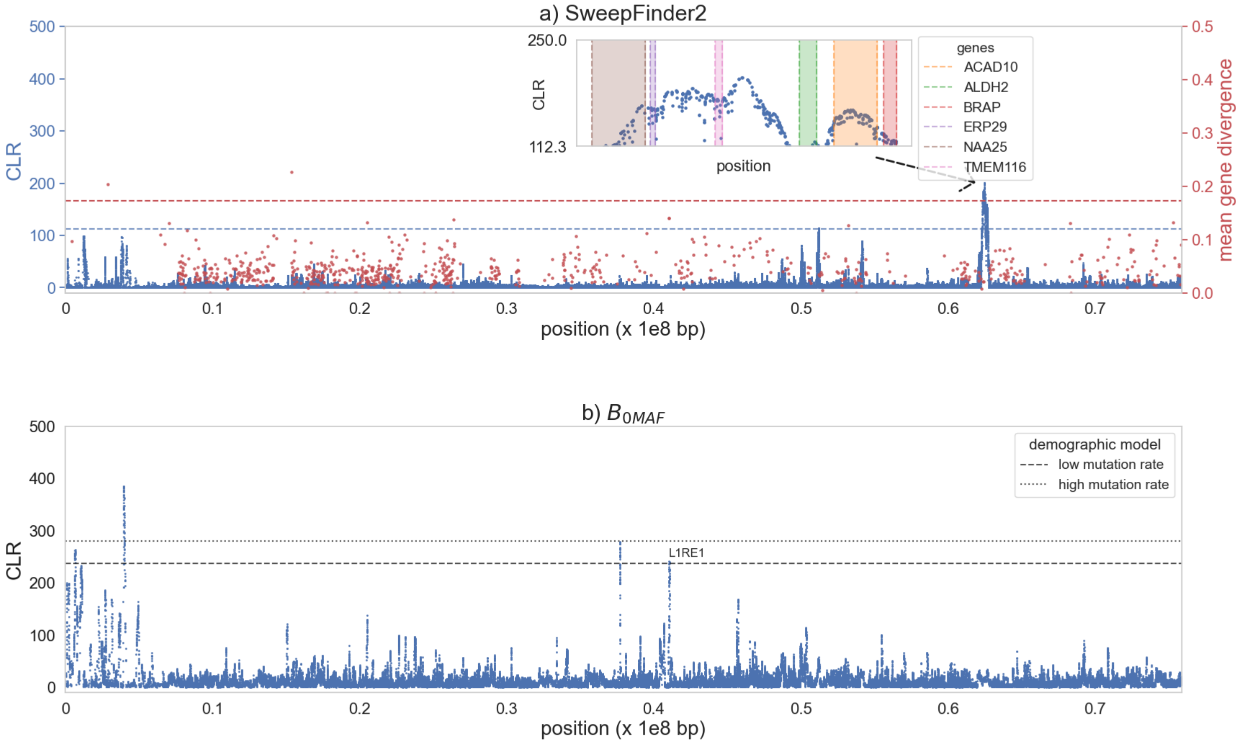


**Supplementary Figure S12: Genome scan results for chromosome 10 using a) SweepFinder2 (shown in blue) and empirical exonic divergence (red); and b) $B_{0MAF}$. a)** The x-axis shows the position along the chromosome, the left y-axis shows the composite likelihood ratio (CLR) value of the sweep statistic at each SNP, and the right y-axis provides the mean gene divergence. The horizontal blue dashed line represents the null threshold for sweep detection, and the horizontal red dashed line represents the 99.9$^{th}$ percentile neutral divergence. Inset plots zoom in on likelihood surface peaks, with genes in these regions highlighted. **b)** The x-axis shows the position along the chromosome, and the y-axis the CLR value of the balancing selection statistic at each fifth SNP. The horizontal dashed lines represent the null thresholds for detection of balancing selection, based on two different coppery titi demographic models (see the “Materials and Methods” section for details). Candidate genes are labeled.

### Supplementary Figure S13

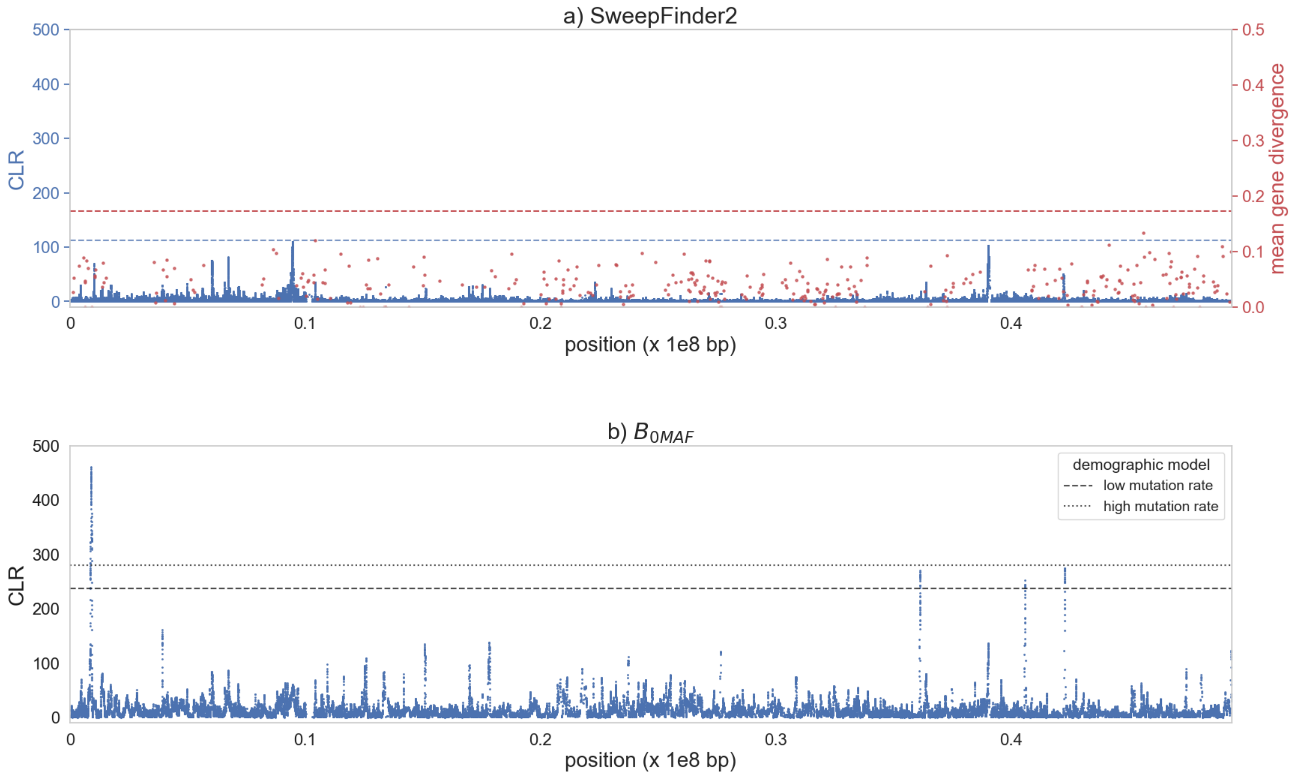


**Supplementary Figure S13: Genome scan results for chromosome 11 using a) SweepFinder2 (shown in blue) and empirical exonic divergence (red); and b)** $\boldsymbol{B_{0MAF}}$**. a)** The x-axis shows the position along the chromosome, the left y-axis shows the composite likelihood ratio (CLR) value of the sweep statistic at each SNP, and the right y-axis provides the mean gene divergence. The horizontal blue dashed line represents the null threshold for sweep detection, and the horizontal red dashed line represents the 99.9$^{th}$ percentile neutral divergence. **b)** The x-axis shows the position along the chromosome, and the y-axis the CLR value of the balancing selection statistic at each fifth SNP. The horizontal dashed lines represent the null thresholds for detection of balancing selection, based on two different coppery titi demographic models (see the "Materials and Methods" section for details).

**Supplementary Figure S14**

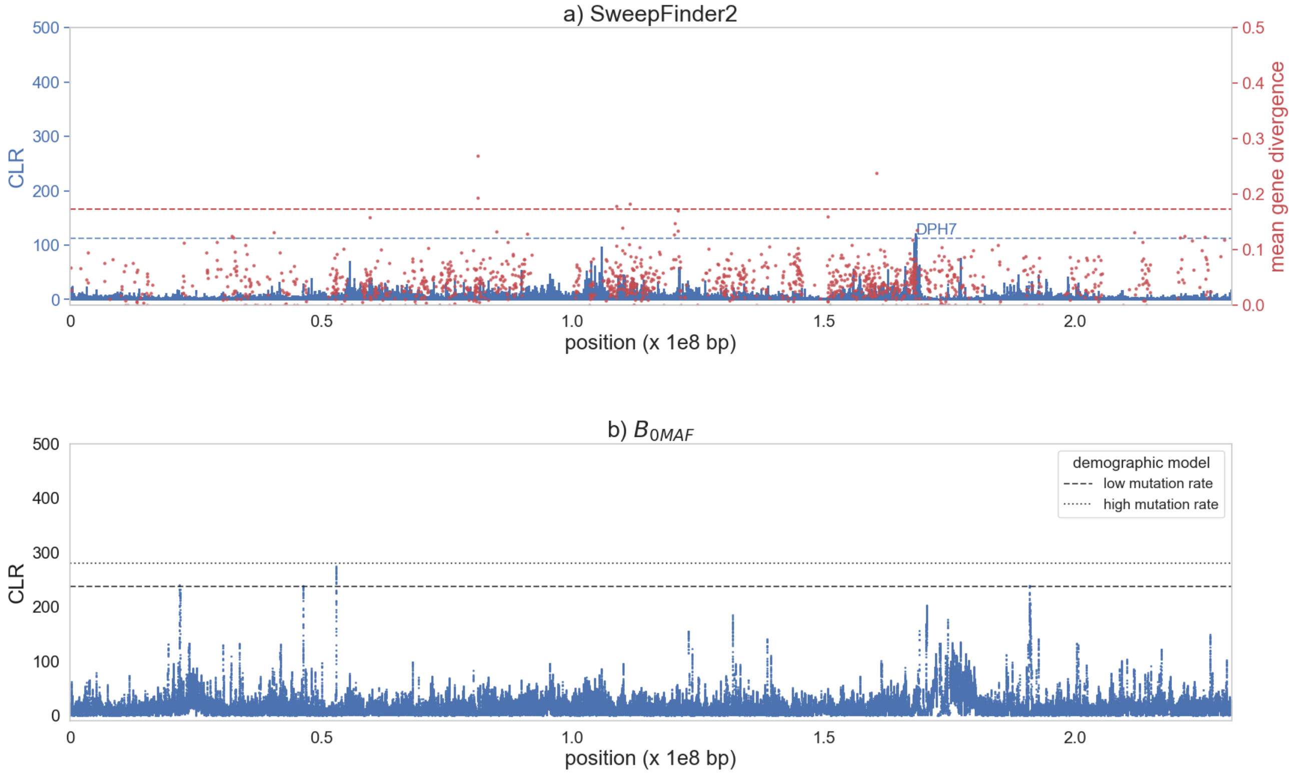


**Supplementary Figure S14: Genome scan results for chromosome 12 using a) SweepFinder2 (shown in blue) and empirical exonic divergence (red); and b) $B_{0MAF}$. a)** The x-axis shows the position along the chromosome, the left y-axis shows the composite likelihood ratio (CLR) value of the sweep statistic at each SNP, and the right y-axis provides the mean gene divergence. The horizontal blue dashed line represents the null threshold for sweep detection, and the horizontal red dashed line represents the 99.9$^{th}$ percentile neutral divergence. **b)** The x-axis shows the position along the chromosome, and the y-axis the CLR value of the balancing selection statistic at each fifth SNP. The horizontal dashed lines represent the null thresholds for detection of balancing selection, based on two different coppery titi demographic models (see the "Materials and Methods" section for details).

**Supplementary Figure S15**

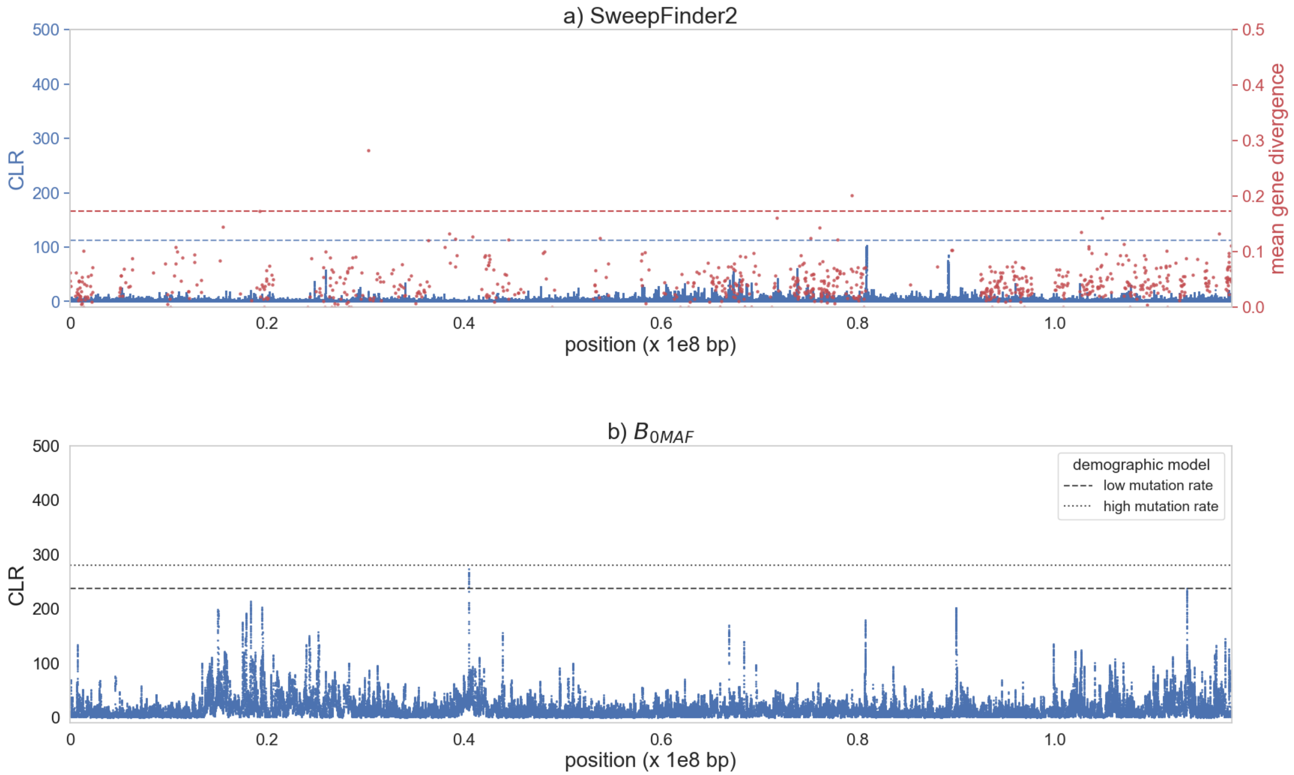


**Supplementary Figure S15: Genome scan results for chromosome 13 using a) SweepFinder2 (shown in blue) and empirical exonic divergence (red); and b) $B_{0MAF}$. a)** The x-axis shows the position along the chromosome, the left y-axis shows the composite likelihood ratio (CLR) value of the sweep statistic at each SNP, and the right y-axis provides the mean gene divergence. The horizontal blue dashed line represents the null threshold for sweep detection, and the horizontal red dashed line represents the 99.9th percentile neutral divergence. **b)** The x-axis shows the position along the chromosome, and the y-axis the CLR value of the balancing selection statistic at each fifth SNP. The horizontal dashed lines represent the null thresholds for detection of balancing selection, based on two different coppery titi demographic models (see the “Materials and Methods” section for details).

**Supplementary Figure S16**

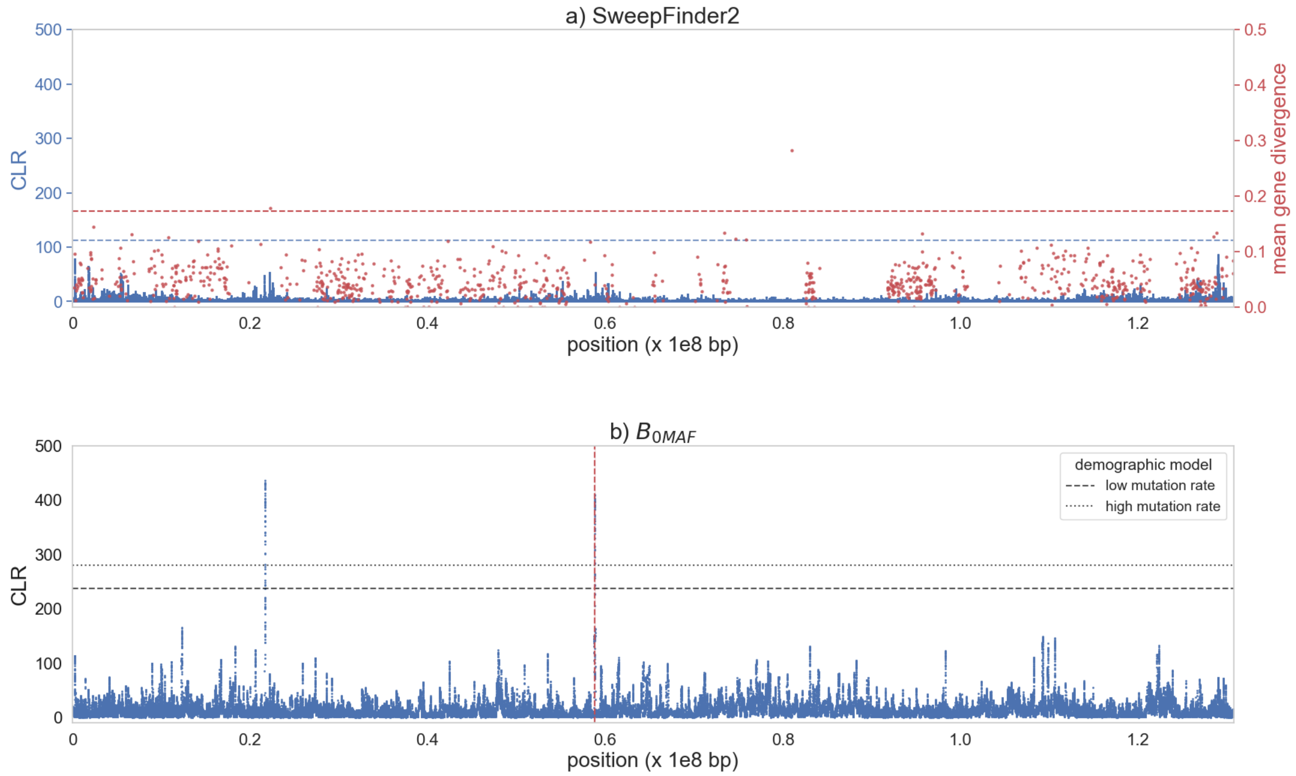


**Supplementary Figure S16: Genome scan results for chromosome 14 using a) SweepFinder2 (shown in blue) and empirical exonic divergence (red); and b) $B_{0MAF}$. a)** The x-axis shows the position along the chromosome, the left y-axis shows the composite likelihood ratio (CLR) value of the sweep statistic at each SNP, and the right y-axis provides the mean gene divergence. The horizontal blue dashed line represents the null threshold for sweep detection, and the horizontal red dashed line represents the 99.9$^{th}$ percentile neutral divergence. **b)** The x-axis shows the position along the chromosome, and the y-axis the CLR value of the balancing selection statistic at each fifth SNP. The horizontal dashed lines represent the null thresholds for detection of balancing selection, based on two different coppery titi demographic models (see the "Materials and Methods" section for details). The position of the deletion is marked by the vertical dashed red line.

**Supplementary Figure S17**

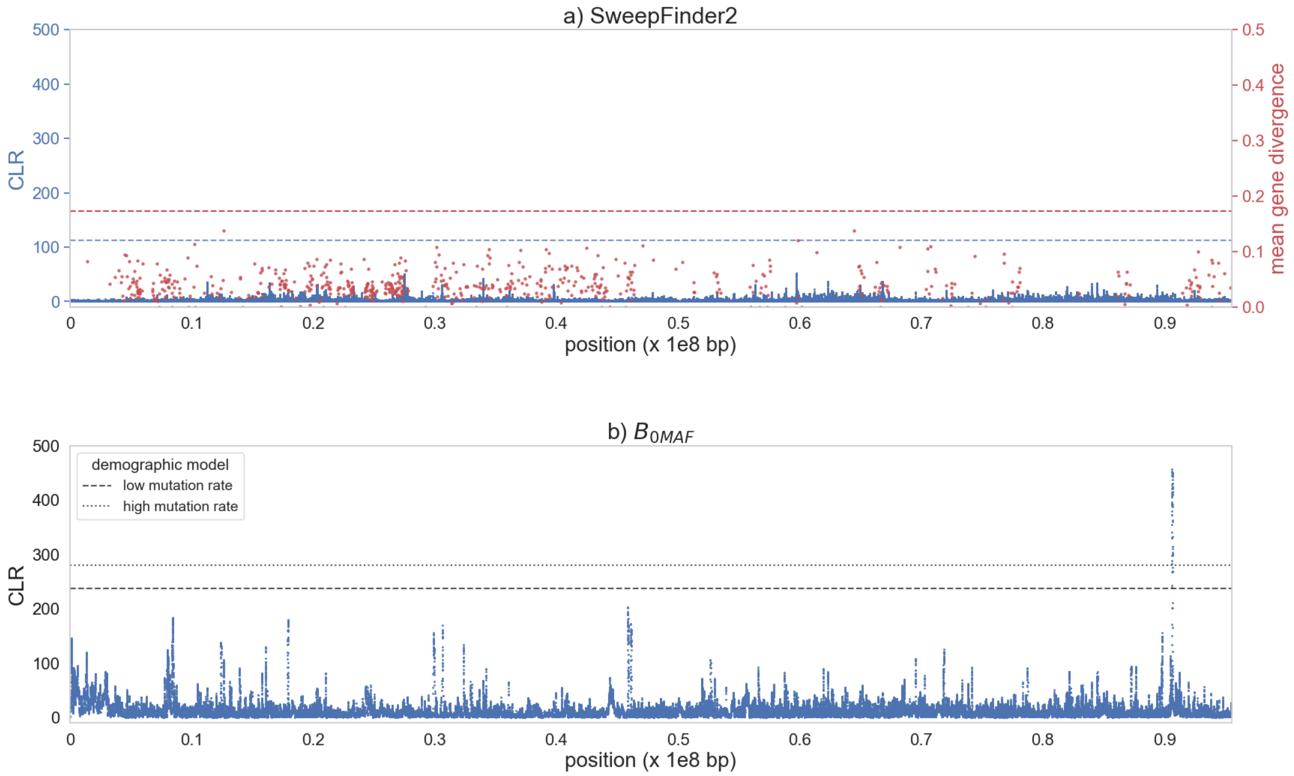


**Supplementary Figure S17: Genome scan results for chromosome 15 using a) SweepFinder2 (shown in blue) and empirical exonic divergence (red); and b) $B_{0MAF}$. a)** The x-axis shows the position along the chromosome, the left y-axis shows the composite likelihood ratio (CLR) value of the sweep statistic at each SNP, and the right y-axis provides the mean gene divergence. The horizontal blue dashed line represents the null threshold for sweep detection, and the horizontal red dashed line represents the 99.9$^{th}$ percentile neutral divergence. **b)** The x-axis shows the position along the chromosome, and the y-axis the CLR value of the balancing selection statistic at each fifth SNP. The horizontal dashed lines represent the null thresholds for detection of balancing selection, based on two different coppery titi demographic models (see the "Materials and Methods" section for details).

**Supplementary Figure S18**

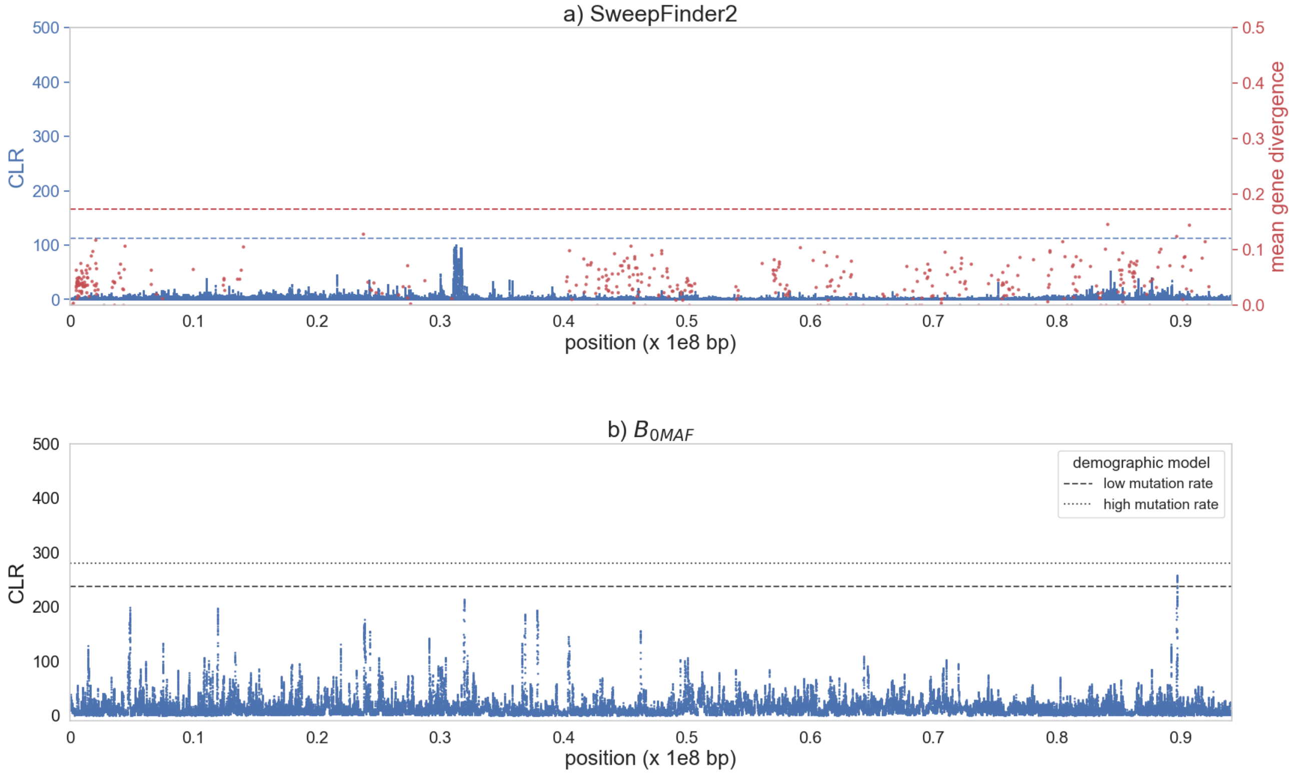


**Supplementary Figure S18: Genome scan results for chromosome 16 using a) SweepFinder2 (shown in blue) and empirical exonic divergence (red); and b) $B_{0MAF}$. a)** The x-axis shows the position along the chromosome, the left y-axis shows the composite likelihood ratio (CLR) value of the sweep statistic at each SNP, and the right y-axis provides the mean gene divergence. The horizontal blue dashed line represents the null threshold for sweep detection, and the horizontal red dashed line represents the 99.9$^{th}$ percentile neutral divergence. **b)** The x-axis shows the position along the chromosome, and the y-axis the CLR value of the balancing selection statistic at each fifth SNP. The horizontal dashed lines represent the null thresholds for detection of balancing selection, based on two different coppery titi demographic models (see the "Materials and Methods" section for details).

**Supplementary Figure S19**

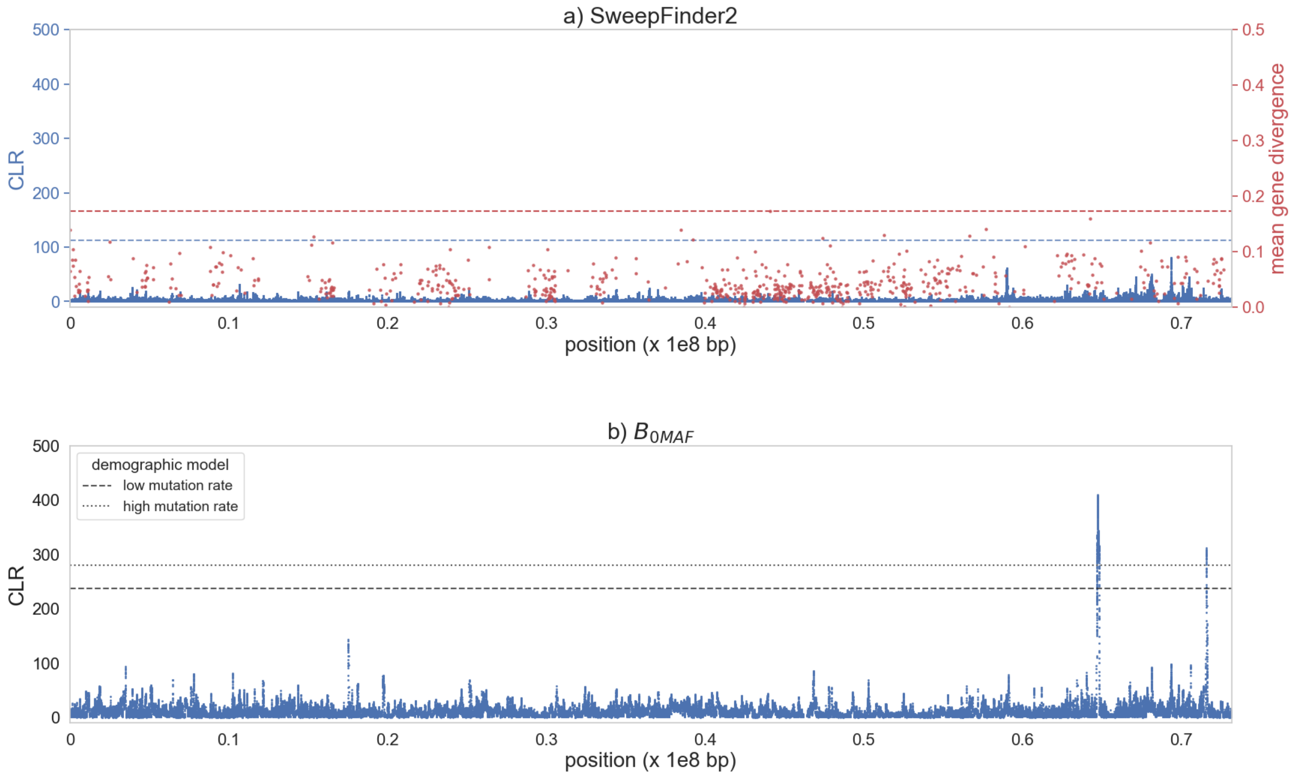


**Supplementary Figure S19: Genome scan results for chromosome 17 using a) SweepFinder2 (shown in blue) and empirical exonic divergence (red); and b) $B_{0MAF}$. a)** The x-axis shows the position along the chromosome, the left y-axis shows the composite likelihood ratio (CLR) value of the sweep statistic at each SNP, and the right y-axis provides the mean gene divergence. The horizontal blue dashed line represents the null threshold for sweep detection, and the horizontal red dashed line represents the 99.9$^{th}$ percentile neutral divergence. **b)** The x-axis shows the position along the chromosome, and the y-axis the CLR value of the balancing selection statistic at each fifth SNP. The horizontal dashed lines represent the null thresholds for detection of balancing selection, based on two different coppery titi demographic models (see the "Materials and Methods" section for details).

**Supplementary Figure S20**

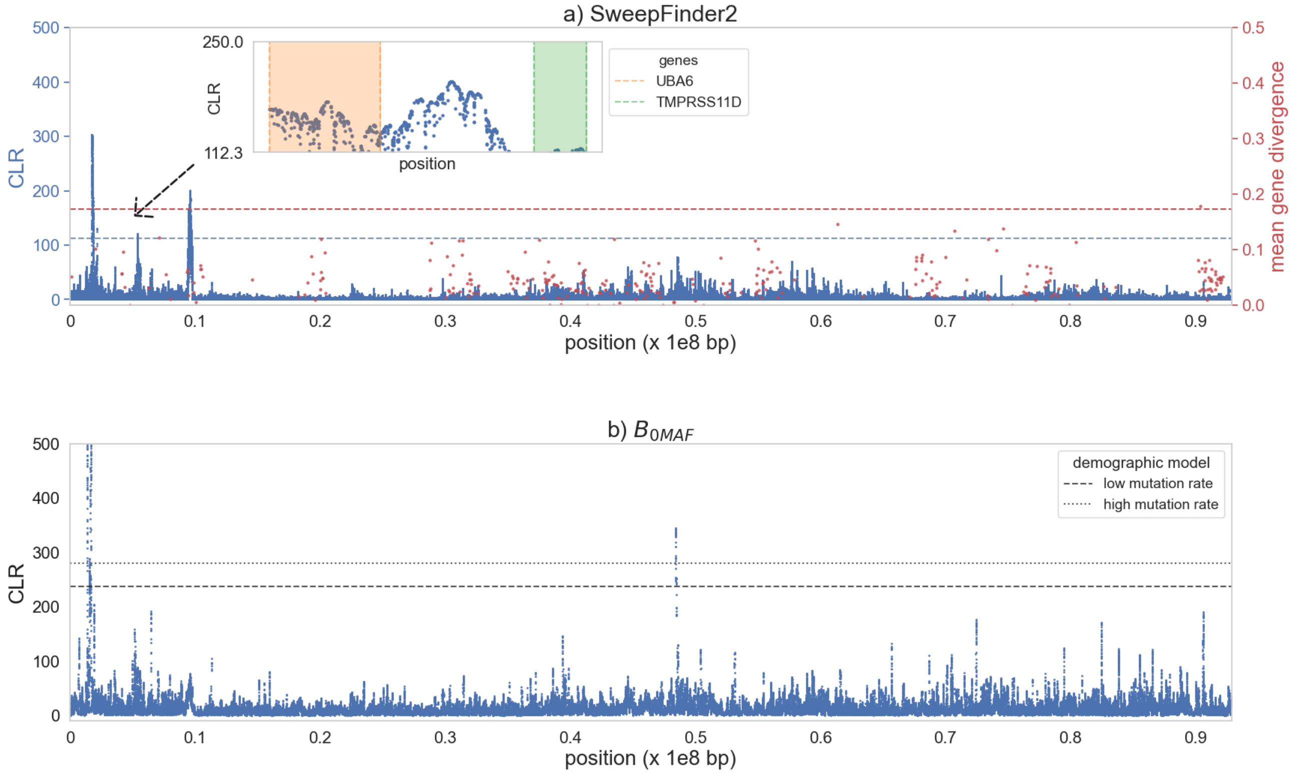


**Supplementary Figure S20: Genome scan results for chromosome 18 using a) SweepFinder2 (shown in blue) and empirical exonic divergence (red); and b) $B_{0MAF}$. a)** The x-axis shows the position along the chromosome, the left y-axis shows the composite likelihood ratio (CLR) value of the sweep statistic at each SNP, and the right y-axis provides the mean gene divergence. The horizontal blue dashed line represents the null threshold for sweep detection, and the horizontal red dashed line represents the 99.9$^{th}$ percentile neutral divergence. Inset plots zoom in on likelihood surface peaks, with genes in these regions highlighted. **b)** The x-axis shows the position along the chromosome, and the y-axis the CLR value of the balancing selection statistic at each fifth SNP. The horizontal dashed lines represent the null thresholds for detection of balancing selection, based on two different coppery titi demographic models (see the "Materials and Methods" section for details).

**Supplementary Figure S21**

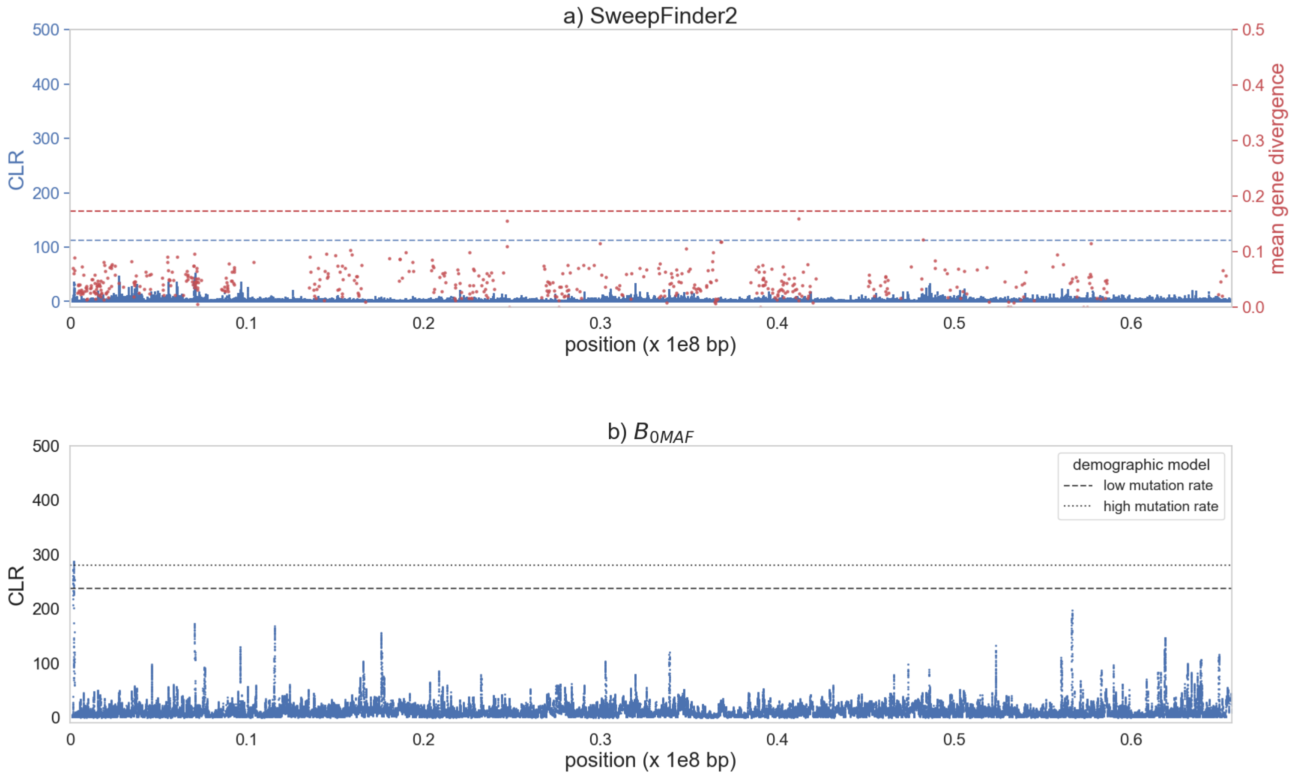


**Supplementary Figure S21: Genome scan results for chromosome 19 using a) SweepFinder2 (shown in blue) and empirical exonic divergence (red); and b) $B_{0MAF}$. a)** The x-axis shows the position along the chromosome, the left y-axis shows the composite likelihood ratio (CLR) value of the sweep statistic at each SNP, and the right y-axis provides the mean gene divergence. The horizontal blue dashed line represents the null threshold for sweep detection, and the horizontal red dashed line represents the 99.9$^{th}$ percentile neutral divergence. **b)** The x-axis shows the position along the chromosome, and the y-axis the CLR value of the balancing selection statistic at each fifth SNP. The horizontal dashed lines represent the null thresholds for detection of balancing selection, based on two different coppery titi demographic models (see the “Materials and Methods” section for details).

**Supplementary Figure S22**

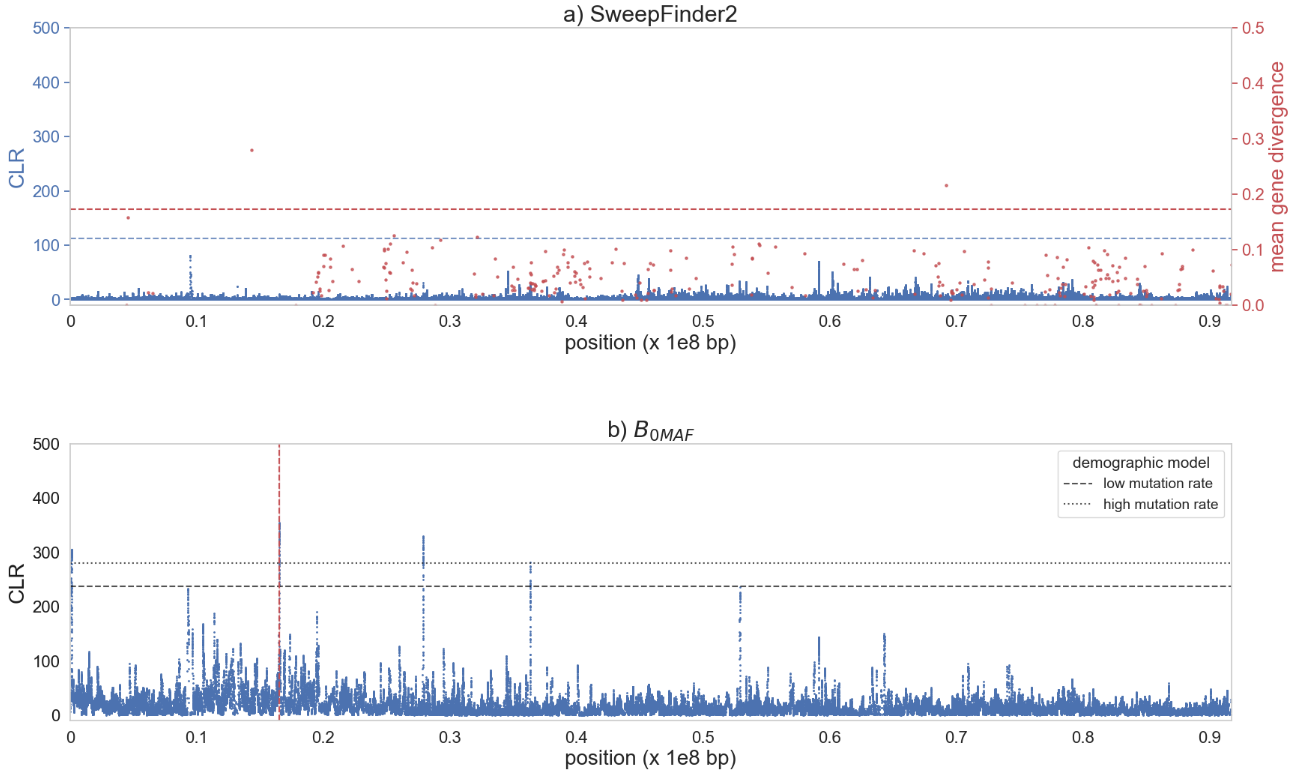


**Supplementary Figure S22: Genome scan results for chromosome 20 using a) SweepFinder2 (shown in blue) and empirical exonic divergence (red); and b) $B_{0MAF}$. a)** The x-axis shows the position along the chromosome, the left y-axis shows the composite likelihood ratio (CLR) value of the sweep statistic at each SNP, and the right y-axis provides the mean gene divergence. The horizontal blue dashed line represents the null threshold for sweep detection, and the horizontal red dashed line represents the 99.9$^{th}$ percentile neutral divergence. **b)** The x-axis shows the position along the chromosome, and the y-axis the CLR value of the balancing selection statistic at each fifth SNP. The horizontal dashed lines represent the null thresholds for detection of balancing selection, based on two different coppery titi demographic models (see the “Materials and Methods” section for details). The position of the deletion is marked by the vertical dashed red line.

**Supplementary Figure S23**

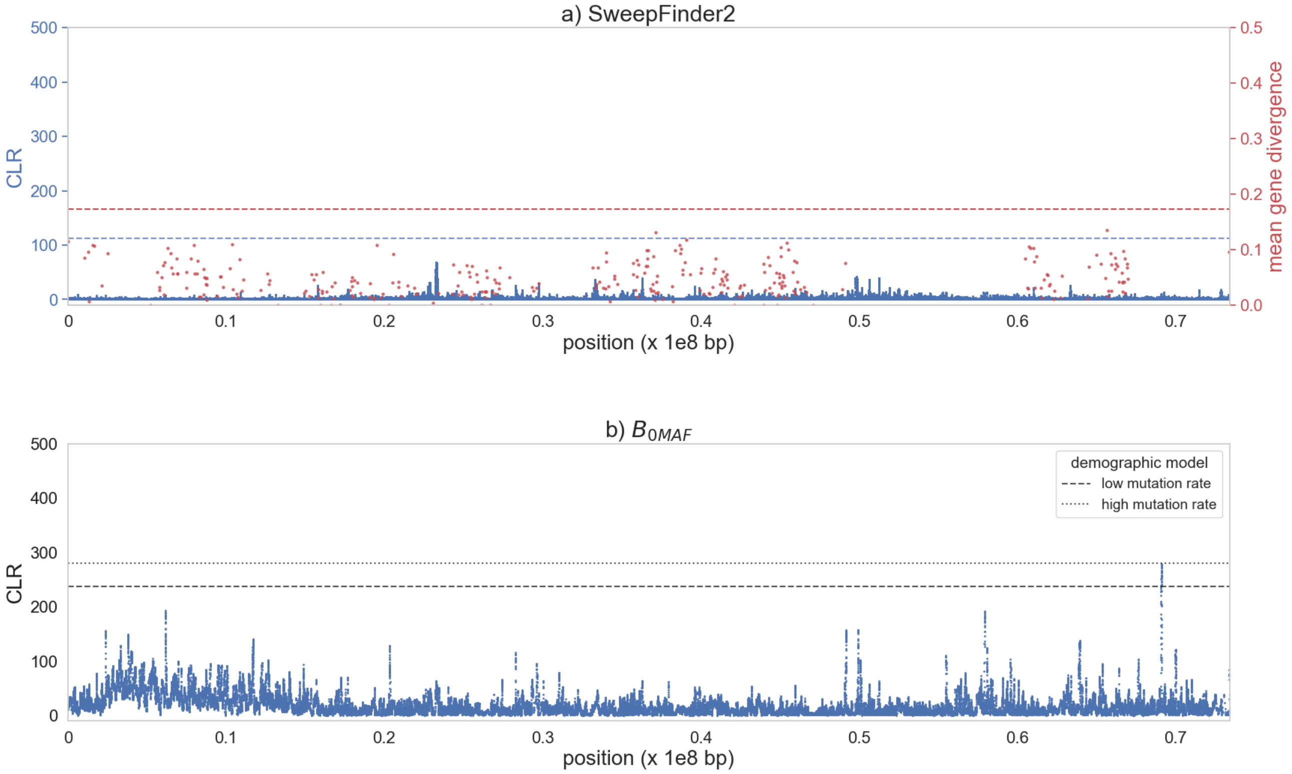


**Supplementary Figure S23: Genome scan results for chromosome 21 using a) SweepFinder2 (shown in blue) and empirical exonic divergence (red); and b) $B_{0MAF}$. a)** The x-axis shows the position along the chromosome, the left y-axis shows the composite likelihood ratio (CLR) value of the sweep statistic at each SNP, and the right y-axis provides the mean gene divergence. The horizontal blue dashed line represents the null threshold for sweep detection, and the horizontal red dashed line represents the 99.9$^{th}$ percentile neutral divergence. **b)** The x-axis shows the position along the chromosome, and the y-axis the CLR value of the balancing selection statistic at each fifth SNP. The horizontal dashed lines represent the null thresholds for detection of balancing selection, based on two different coppery titi demographic models (see the "Materials and Methods" section for details).

**Supplementary Figure S24**

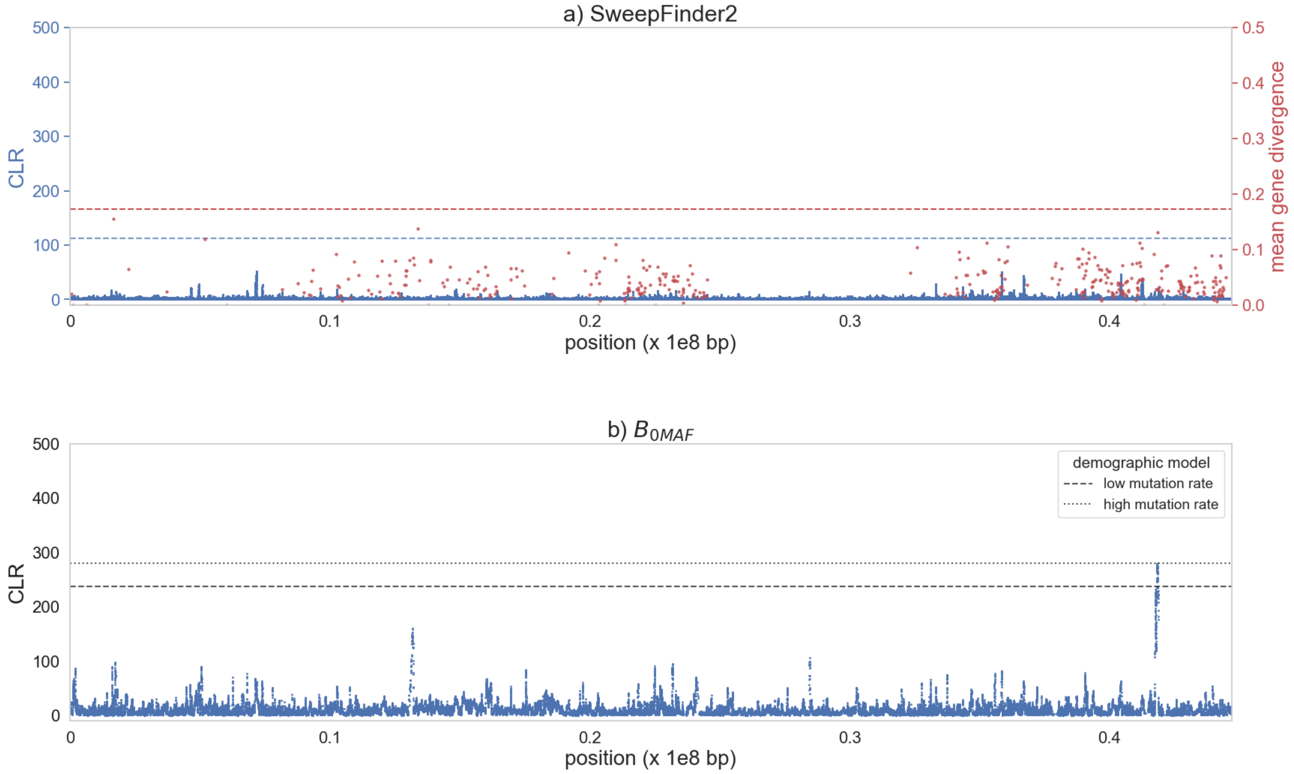


**Supplementary Figure S24: Genome scan results for chromosome 22 using a) SweepFinder2 (shown in blue) and empirical exonic divergence (red); and b) $B_{0MAF}$. a)** The x-axis shows the position along the chromosome, the left y-axis shows the composite likelihood ratio (CLR) value of the sweep statistic at each SNP, and the right y-axis provides the mean gene divergence. The horizontal blue dashed line represents the null threshold for sweep detection, and the horizontal red dashed line represents the 99.9$^{th}$ percentile neutral divergence. **b)** The x-axis shows the position along the chromosome, and the y-axis the CLR value of the balancing selection statistic at each fifth SNP. The horizontal dashed lines represent the null thresholds for detection of balancing selection, based on two different coppery titi demographic models (see the "Materials and Methods" section for details).
